# Passaging Human Tauopathy Patient Samples in Cells Generates Heterogeneous Fibrils with a Subpopulation Adopting Disease Folds

**DOI:** 10.1101/2023.07.19.549721

**Authors:** Zhikai Zeng, Karen Tsay, Vishnu Vijayan, Matthew P. Frost, Shikhar Prakash, Athena Quddus, Alexa Albert, Michael Vigers, Madhur Srivastava, Amanda L. Woerman, Songi Han

## Abstract

The recent discovery by cryo-electron microscopy (cryo-EM) that the neuropathological hallmarks of different tauopathies, including Alzheimer’s disease, corticobasal degeneration (CBD), and progressive supranuclear palsy (PSP), are caused by unique misfolded conformations of the protein tau is among the most profound developments in neurodegenerative disease research. To capitalize on these discoveries for therapeutic development, one must achieve *in vitro* replication of tau fibrils that adopt the representative tauopathy disease folds, which represents a grand challenge for the field. An widely used approach is seeded propagation using pathological tau fibrils derived from post-mortem patients in biosensor cells that expresses a fragment of the tau protein, known as K18 or Tau4RD, containing mainly the microtubule-binding repeat domain of tau as the substrate. The new insight from cryo-EM raised the question whether the Tau4RD fragment is capable of adopting the specific tau folds found in CBD and PSP patient fibrils, and whether cell-passaged and amplified tau fibrils can be used as seeds to achieve cell-free assembly of recombinant 4R tau into fibrils without the addition of any cofactors. Using Double Electron Electron Resonance (DEER) spectroscopy, we discovered that cell-passaged pathological seeds generate heterogeneous fibrils that are distinct between the CBD and PSP lysate-seeded fibrils. These fibrils are also distinct from heparin-induced tau fibril populations. Moreover, the lysate-seeded fibrils contain a characteristic sub-population that resembles the disease fold corresponding to the respective starting patient sample. These findings indicate that templated propagation using CBD and PSP patient-derived fibrils is possible with a tau fragment that does not contain the entire pathological fibril core, and that additional mechanisms must be tuned to converge to a homogeneous fibril population.

## 1 Introduction

Tauopathies are a varied group of neurodegenerative diseases defined by the deposition of fibrillar aggregates of the microtubule-binding protein tau (MAPT, or tau) in the brain. Notably, the prevalence of these deposits is closely related to clinically observable neurodegeneration [1, 2]. In the disease state, tau misfolds from a soluble monomer into an insoluble fibril with a *β*-sheet-rich structure capable of recruiting naïve tau to self-replicate, consistent with the prion mechanism of disease. Although tau pathologies specific to each disease vary in morphology and isoform composition [3], the defining difference between the clinical phenotypes is thought to arise from the shape, or conformation, that tau adopts. This is the basis of the strain hypothesis, or the idea that protein conformation determines disease [4–6].

In 2013, the Goedert lab showed that brain homogenates prepared from a variety of deceased tauopathy patient samples, including progressive supranuclear palsy (PSP), corticobasal degeneration (CBD), and argyrophilic grain disease (AGD), induced tau pathology following intracerebral injection into a transgenic mouse model expressing human tau [7]. The induced inclusions exhibited immunostaining patterns similar to those seen in the corresponding human pathology, suggesting that each disease may be caused by a distinct tau strain [7, 8]. Subsequent work investigating tau strain biology employed cell models expressing the microtubule-binding repeat domain of tau, including the fragment spanning residues 244-372 known as K18, with a C-terminal fluorescent reporter protein [9–11]. These studies reported that tau strains are distinct with respect to the seeding capacity, isoform involvement, cell-type specificity, and protease digestion or guanidine denaturation patterns [9–16]. However, it was advances in cryo-electron microscopy (cryo-EM) that facilitating a major leap in our understanding of tau strains on a molecular level, resulting in the resolution of misfolded tau folds present in several tauopathies, including Alzheimer’s disease (AD) [17], Pick’s disease, [18],CBD [19, 20], chronic traumatic encephalopathy (CTE) [21], PSP [22, 23], AGD [23], and globular glial tauopathy (GGT) [23] patient samples. This important body of work has shown that underlying each tauopathy is a distinct conformation of misfolded tau.

Recombinant tau fibrils induced by polyanionic cofactors, such as heparin, were commonly used to study tau aggregation [24], but cryo-EM [25] and Double Electron Electron Resonance (DEER) [26] have revealed that cofactor-induced fibrils are heterogeneous and distinct from patient-derived fibrils, and are thus not disease relevant. This illustrated a need for developing accessible and reliable methods for replicating disease folds. Furthermore, cell-free methods are desirable for structural biology techniques. Previous studies have shown that patient-derived fibrils can propagate in cell-free or *in vitro* systems, as evidenced by the similar morphological and biochemical properties between the tau seeds and resulting fibrils [13, 14, 27–33]. However, whether or not the atomic conformational features of the disease folds are conserved and replicated by second generation seeding has not been investigated to date. We, therefore, set out to test the widely assumed hypothesis that patient-derived fibrils induce tauopathy-specific protein misfolding of recombinant tau monomer after passaging through Tau4RD (residues 244-378) reporter cells. While cryo-EM and solid-state NMR have been crucial for unraveling the atomic basis of tau strains at high-resolution, these are not suitable tools for capturing dynamically evolving and long-range intra-tau distances, nor for studying a heterogeneous fibril population. In order to study *whether* and *how* tau fibrils template protein misfolding in a strain-specific manner at sufficiently high-resolution to differentiate between key disease folds, a structural biology tool is needed that can capture conformations of disordered and partially disordered fibril regions and report on the distribution of heterogeneous fibril populations. Double Electron Electron Resonance (DEER), a pulsed dipolar electron paramagnetic resonance (EPR) spectroscopic method, is such a tool capable of reporting on the distribution of conformations of doubly spin-labeled proteins regardless of its order and disorder, and hence capture the complete population of tau fibrils formed upon seeded aggregation using CBD and PSP patient-derived materials.

In this study, we isolated tau fibrils from post-mortem CBD and PSP patient samples, which we used to establish monoclonal Tau4RD*LM-YFP cell lines expressing Tau4RD (residues 244-378 with the P301L and V337M mutations), also referred to as biosensor cell lines, that stably propagate either CBD or PSP patient-derived fibrils, as shown in Fig. 1a. Lysates collected from these cells were then added to recombinant 4R tau monomer, either Tau187 (residues 255-441) or the 0N4R tau isoform, to seed protein misfolding. DEER was used to measure the probability distribution, P(*r*), of intra-molecular distances *r*, from 1.5 to several nm ranges across a pair of spin-labels, both attached to the same tau molecule. We carefully selected three spin-label pairs to capture distances across tau segments specific to the CBD and PSP folds - residues 351 & 373, 334 & 360, and 340 & 378 (illustrated in Fig. 1b). A similar approach of relying on select spin label pairs on tau molecules to evaluate the protein fold of tau stacked into fibrils was demonstrated in earlier studies [26, 34, 35]. The theoretical distance distribution for the above-listed spin-label pairs on the reported CBD and PSP folds is computed with the RotamerConvolveMD method [36] (shown in Fig. 1c). The doubly spin-labeled tau was diluted with unlabeled, cysteine-less tau to capture intra-molecular distances that are sensitive to the fold of the tau protein stacked into fibrils, and not to inter-molecular distances that are sensitive to the fibril packing order. We compared the measured P(*r*) with the simulated P(*r*) based on the cryo-EM structures of CBD and PSP tau to understand whether fibrils seeded with cell-passaged CBD and PSP tau fibrils are conformationally distinct from each other, and from heparin-induced fibrils. We validated this comparison using the novel frequency pattern analysis, both qualitatively by visualizing the localized frequency differences, and quantitatively by calculating the structure similarity index measure (SSIM) between differently generated fibrils. Moreover, we set out to discover whether a sub-population of the heterogeneous fibril mixture represents inter-molecular tau distances contained in the CBD and PSP disease folds. Given that 1) CBD and PSP patient-derived fibrils are used to induce disease-relevant protein misfolding of Tau4RD that is shorter than the sequence making up the core of the CBD and PSP fibril, 2) no co-factors were used, 3) no post-translational modifications were included, and 4) the experiments were performed in a cell-free system, the answers to the posed questions were not obvious. Thus, this study provides a novel framework for probing disease-specific tau fibril structures and uncovers distinct conformational characteristics that deepen our understanding of the molecular underpinnings of tauopathies like CBD and PSP.

**Fig. 1.**
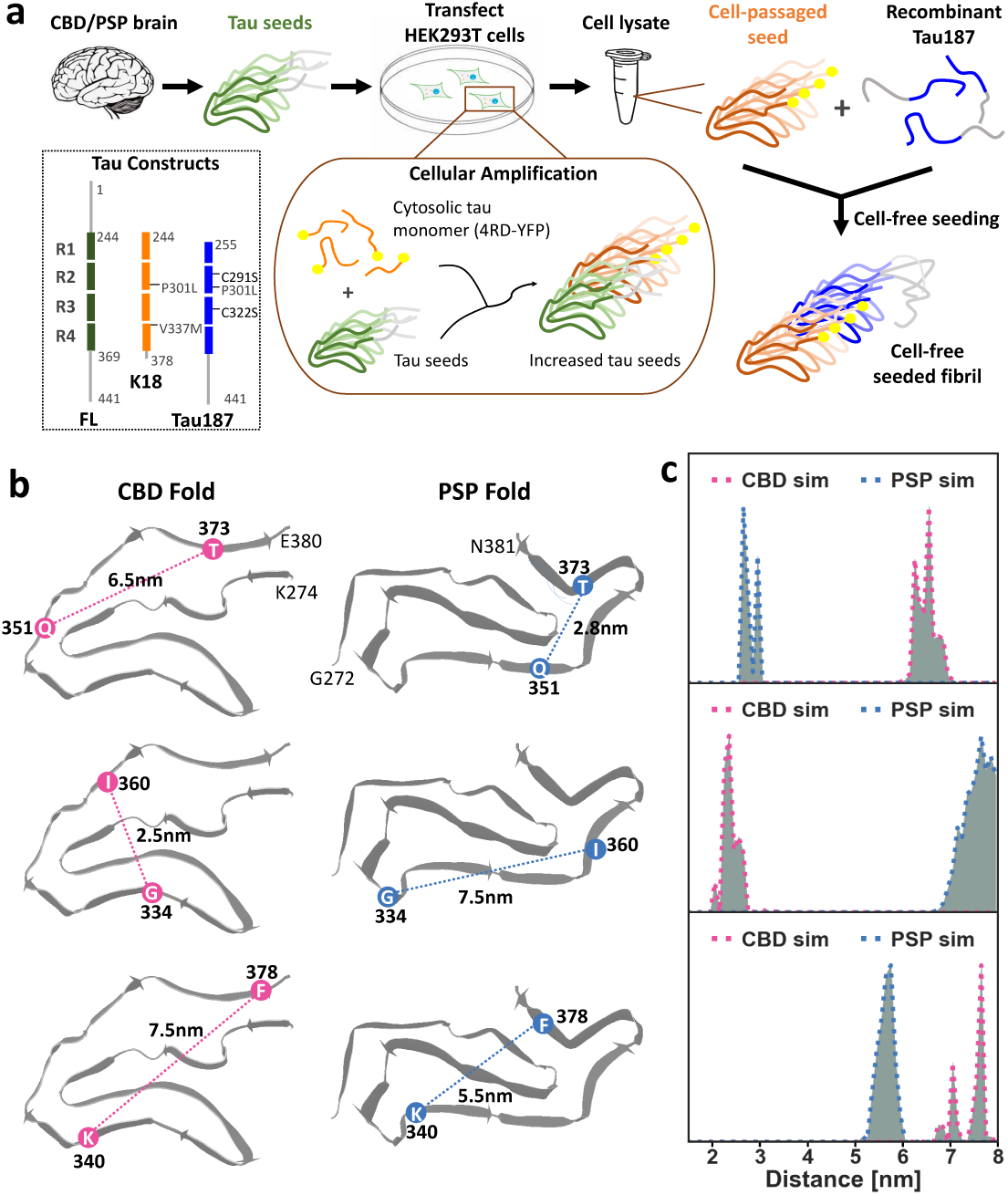
(a) Schematic of the cellular amplification of CBD and PSP patient samples followed by cell-free seeding of recombinant Tau187 using cell-passaged seeds. The inset shows the tau constructs used, including full-length 2N4R tau (FL, in green), Tau4RD (4RD, in orange), and Tau187 (in blue). (b) Schematics of the published CBD fold (PDB ID 6TJO) [21] and PSP fold (PDB ID 7P65) [19] with the three selected spin-label pairs: 351 & 373, 334 & 360, and 340 & 378. The mean distances between the spin-label pairs were measured directly in the PDB files, and are labeled within this figure. (c) Theoretical distance distributions simulated from the published cryo-EM structures of the CBD fold [21] and PSP fold [19] by the RotamerConvolveMD method [36] in the 2-8 nm range for the three spin-labeled pairs. The distance distributions across the spin-label pair at residues 351 & 373 are referred to as P(*r*, 351-373). The same rule applies to P(*r*, 334-360) and P(*r*, 340-378)

## 2 Results

### 2.1 Cell-passaged tauopathy patient samples template recombinant tau misfolding in cell-free system

To investigate tau misfolding in CBD and PSP, we first established Tau4RD*LM-YFP cells that stably propagate pathogenic tau isolated from brain samples received from deceased CBD and PSP patients (schematic shown in Fig. 1a). Using similar methods as previously described [11], tau prions were isolated from one control, one CBD, and one PSP brain homogenate with sodium phosphotungstate (PTA) that selectively precipitates insoluble protein aggregates [37, 38]. The resulting protein pellets were then resuspended in DPBS and incubated with either Tau3RD*VM-YFP cells or Tau4RD*LM-YFP cells for 4 days. Consistent with our previous findings [37], the CBD and PSP patient samples induced tau misfolding and aggregation in the Tau4RD*LM-YFP cells, but had no effect on the Tau3RD*VM-YFP cell line (Fig. 2a & b). We then generated monoclonal cell lines that stably propagate either CBD or PSP tau via single-cell plating of infected cells. Expression levels of the 4RD-YFP construct were confirmed via Western blot using lysate collected in RIPA buffer (Fig. 2c).

**Fig. 2.**
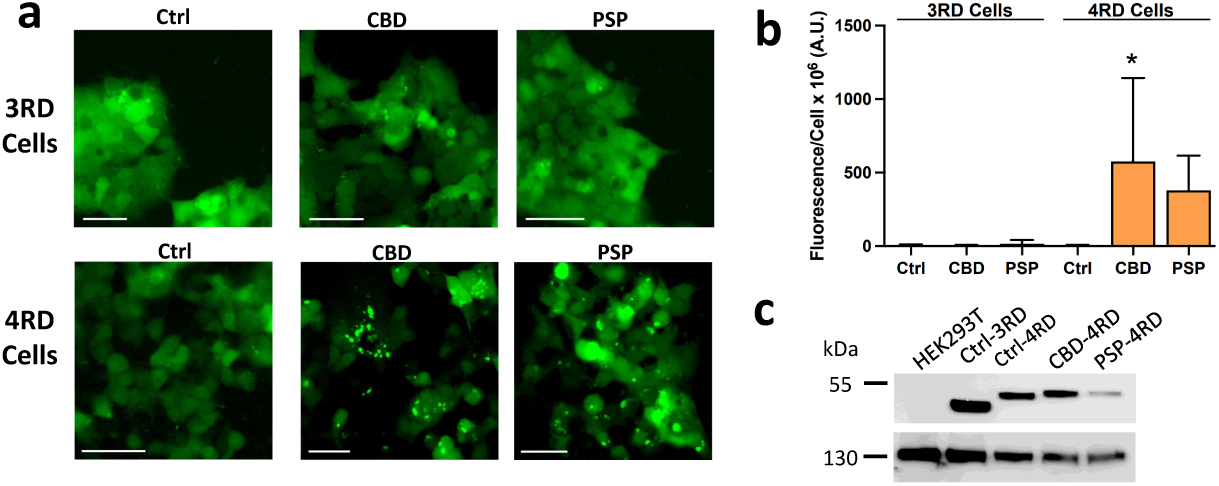
(a) Representative images of Tau4RD*LM–YFP cells and Tau3RD*VM-YFP cells infected with CBD and PSP patient samples. YFP is shown in green. (Scale bar, 50 *µ*m). (b) Quantification of cell infection. * = *P* < 0.05. (c) Lysates collected in RIPA buffer from naïve HEK293T cells, Tau3RD*VM-YFP cells, Tau4RD*LM-YFP cells, and Tau4RD*LM-YFP cells stably infected with CBD or PSP tau prions were analyzed for the presence of the tau-YFP fusion protein by Western blot using the GFP primary antibody (top blot). Vinculin (bottom blot) is shown as a loading control. Full uncropped gel is shown in Fig. S1.

For seeding experiments, lysates from Tau4RD*LM-YFP cells infected with CBD and PSP patient samples were collected in 1X protease inhibitor and further diluted in 20 mM HEPES buffer (pH 7.4) to 5-15% (protein mass percentage) to seed the fibrillization of recombinant Tau187. The two naturally occurring cysteines in this construct were mutated to serines (C291S and C322S) to avoid the formation of disulfide bonds that are known to not be part of pathological tau fibril folds [19, 21], and instead may inhibit tau aggregation [39]. To determine if the cell-passaged lysates can induce the fibrillization of tau, thioflavin T (ThT) fluorescence measurements were collected over the course of 24 hours. ThT is a fluorescent dye that binds specifically to *β*-sheet structures of amyloid proteins [40] and provides an *in situ* assessment of *β*-sheet abundance. The cell-passaged CBD and PSP lysates induced a robust increase in ThT fluorescence of Tau187 in a concentration-dependent manner (fluorescence intensity increased when 15% lysate was used compared to 5%; Fig. 3a). Not surprisingly, when polydisperse heparin (average molecular mass 15 kDa, Galen Lab Inc.) was added to the recombinant tau at a tau:heparin molar ratio of 4:1 (equivalent to 17% heparin by mass), heparin induced greater fibril quantities than the CBD or PSP lysates, as assessed by a significantly greater increase in the ThT max fluorescence emission (*** = *P* < 0.001). The negative control, lysate from uninfected Tau4RD cells, did not induce fibrillization (Fig. 3a). Furthermore, the cell-passaged lysates were also used to successfully seed the fibrillization of recombinant full-length (FL) 0N4R tau, but again less than what heparin induced as determined via ThT fluorescence (Fig. S2a). We focused on Tau187 in the subsequent experiments as it has a more robust aggregation propensity compared to FL 0N4R tau. Tau187 contains the entire Tau4RD sequence, as well as all regions included in the CBD and PSP fibril cores, and hence is ideally suited to study the seeded aggregation by cell-passaged CBD or PSP lysates.

**Fig. 3.**
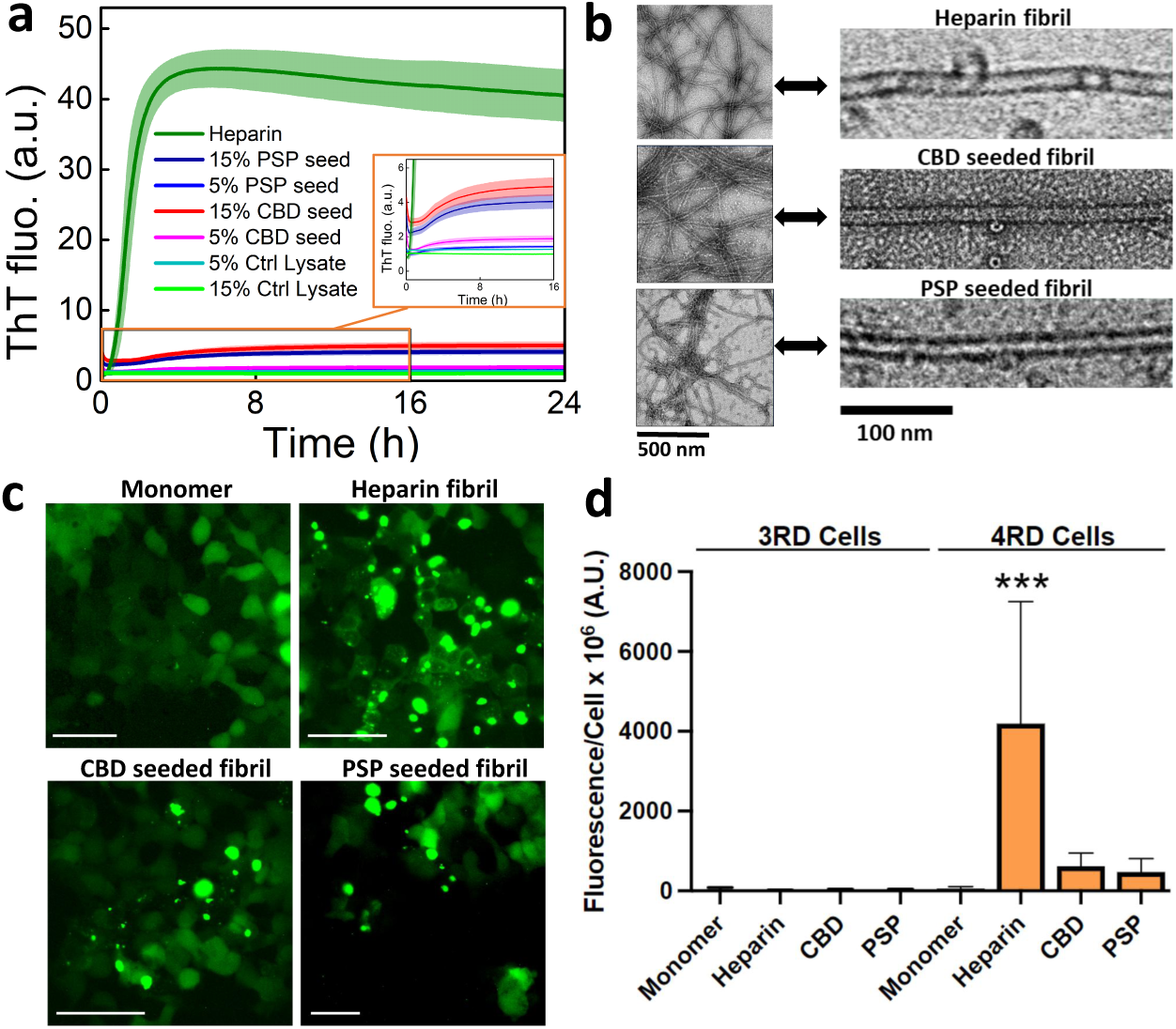
(a) ThT fluorescence of tau187 fibrillization induced by heparin (green), Tau4RD cell-passaged seeds (lysates) from CBD (pink) or PSP (blue) patient samples for Tau187 construct, or uninfected Tau4RD cell lysate (negative control, cyan and light green). The CBD and PSP lysate-seeded aggregation generated a relatively low maximum ThT while the control lysate did not generate any ThT signal increase; an enlarged curve is shown in the inset. Tau187 (25 *µ*M) was mixed with stoichiometric amounts of heparin (8.25 *µ*M, 17% by mass) or cell-passaged lysates [either 5% (lighter-color lines) or 15% by mass (darker-colored lines)] and was aggregated in the presence of ThT at 37 °C. (b) Representative negative stain TEM of heparin-induced fibrils and lysate-seeded fibrils (scale bar, 500 nm.), with magnified images shown to the right (scale bar, 100 nm). Full images are shown in Fig. S3. (c) Representative images of Tau4RD*LM–YFP cells infected with monomeric Tau187, heparin-induced fibrils, and lysate-seeded fibrils. While all fibrils induced tau-YFP aggregation, monomeric Tau187 had no effect on the cells. YFP is shown in green. (Scale bar, 50 *µ*m). (d) Quantification of tau-YFP aggregation following infection with Tau187 monomer, heparin-induced fibrils, or lysate-seeded fibrils in Tau3RD*VM-YFP cells (left) and Tau4RD*LM-YFP cells (right). *** = *P* < 0.001

To ensure that the measured fluorescence increase in the ThT assay was due to the formation of tau fibrils rather than soluble molecules in the cell-passaged seeds, we pelleted the tau aggregates from the lysates by ultracentrifugation at 100,000 x *g* for 1 hour at 4 °C, removing any soluble small molecule species in the supernatant. The remaining tau fibrils were then resuspended in 20 mM HEPES buffer (pH 7.4) and used to seed recombinant Tau187 fibrillization. ThT fluorescence showed that the pelleted samples induced fibrillization, whereas the supernatant did not (Fig. S2b). Moreover, the isolated fibrils and clarified lysates induced comparable fluorescence plateaus, indicating that the seeding capacity of the cell-passaged fibrils originates from the aggregated insoluble tau, not soluble species in the lysates. We can conclude that the cell-passaged fibrils induce recombinant tau misfolding and fibrillization, and the kinetics of this process are influenced by the quantity of the seed used. To maximize the seeded fibril quantity, 15% cell-passaged seeds were used in subsequent experiments.

The morphologies of the heparin-induced and lysate-seeded fibrils were characterized by negative stain Transmission Electron Microscopy (TEM). As shown in Fig. 3b, heparin-induced fibrils were predominantly long, straight, and well-separated from one another with a few of the fibrils exhibiting a wavy appearance. We also observed long and straight, but more associated, fibrils for both CBD and PSP lysate-seeded fibrils. The CBD lysate-seeded fibrils have a diameter of 20 nm and are devoid of any helical twists. Earlier reports suggest that CBD fibrils are heterogeneous, containing both narrow, straight fibrils and wider twisted, ribbon-like fibrils [41–43]. The morphology we observe in this study is closer to the narrow, straight filaments reported for CBD [19]. The TEM images of PSP-induced fibrils showed fibrils with a very similar appearance, featuring slightly smaller diameter fibrils (16-18 nm) with the appearance of straight tubules, in agreement with the previous reports of brain-derived PSP fibril morphology.[42, 43]

To determine if the heparin-induced and lysate-seeded fibrils exhibit prion activity, we incubated both the Tau3RD*VM-YFP and Tau4RD*LM-YFP cells with all three fibril types, as well as monomeric Tau187. Indeed, all three fibrils induced tau aggregation in the Tau4RD*LM-YFP cells but had no effect on the Tau3RD*VM-YFP cells (Fig. 3c & d), consistent with these being 4R fibrils. In contrast, monomeric Tau187 had no effect in either cell line tested.

### 2.2 Tau fibrils seeded using cell-passaged patient samples have distinct conformations with sub-populations that recapitulate disease-specific folds

Although cryo-EM tau structures [19–23, 23, 44] have rightfully captured the field’s attention, cryo-EM can only resolve a homogeneous fibril population from brain samples after extensive purification of fibrils from large quantities of brain material. Moreover, only large, detergent-insoluble fibrils have been successfully imaged, which may not include critical intermediates en route to pathological tau aggregation. In contrast, DEER spectroscopy provides an efficient and direct approach to characterize partially ordered and/or heterogeneous fibrils, or even of disordered proteins and partially aggregated proteins. It resolves the probability distribution of intra-molecular distances in the 1.5 to 8 nm ranges across a pair of spin-labels attached to a single tau molecule. Therefore, DEER can be used for discovery work on heterogeneous samples or to capture transient conformational ensembles populated along the pathway of seeded aggregation. After the conditions for structural convergence are achieved, guided by DEER studies, cryo-EM or other higher-resolution structural biology tools can be used to characterize the end product.

Consistently distinct fibril morphologies observed by negative stain TEM may have different underlying protein folds. However, there are many examples where similar protein folds give rise to distinct fibril morphologies, presumably due to differences in quaternary packing of protofibrils along the fibril axis or inter-fibril interactions, to name a few factors [17, 19, 45]. Some studies have shown different protein folds give rise to indistinguishable fibril morphologies [20], and vice versa. Hence, protein folds within a fibril cannot be derived from ultrastructural properties, but must be directly measured.

To investigate the conformational features of the lysate-seeded fibrils, we employed DEER to measure the distribution of intra-tau distances. DEER has the unique ability to measure the distribution of distances between a pair of electron spin-labels covalently tethered to tau at two sites modified by site-directed mutagenesis to cysteines. DEER is also solely sensitive to paramagnetic probes such as MTSL [S-(1-oxyl-2,2,5,5-tetramethyl-2,5-dihydro-1H-pyrrol-3-yl)methyl methanesulfonothioate] and is not affected by the disorder or size of the protein, and hence it is particularly well suited for tracking protein aggregation. In contrast, the effects of protein crowding, sample heterogeneity, and protein intrinsic disorder render the use of NMR, crystallography, or cryo-EM challenging or infeasible. Previously, we presented DEER-derived distance distributions from pairwise spin-labeled tau stacked to fibrils induced by the addition of the co-factor heparin and found that these fibrils are structurally heterogeneous, judging by the width of the intra-tau distance distribution, P(*r*). These structures were found to be clearly distinct from the tau fibrils present in AD patient samples by comparing the theoretical distance distribution for spin-label pairs on the reported AD fold with the experimentally measured one [26]. This result was subsequently confirmed with cryo-EM structures of heparin-induced tau fibrils that revealed a complete lack of homology between the heparin-induced and patient-derived fibrils [25].

In this study, we followed a similar procedure as reported in Fichou *et al.* [26], rationally selecting the spin-label pairs to be positioned along the outer layer of the fibril cores of the CBD and PSP folds to differentiate between the two cryo-EM structures [19, 21]. Critically, we selected spin-label pairs that yield significantly different P(*r*) between the monomeric and aggregated state (Fig. S4) to ensure that the change of the mean distance and distance distribution can unambiguously report on tau misfolding. All labeled sites were chosen due to their location outside of the heparin-induced fibril core as identified by cryo-EM [25] and hence should differentiate between heparin-induced and patient-derived structures.

After selecting our spin-label pairs, we introduced two cysteines into Tau187 at these desired sites by site-directed mutagenesis and spin-labeling (SDSL) of MTSL to the cysteine residues [46, 47]. To ensure that DEER measurements capture intra-molecular (not inter-molecular) distances, we diluted the doubly spin-labeled Tau187 with unlabeled Tau187 (i.e., without mutation to cysteines, and hence also no labels) to get spin-diluted Tau187 (10% doubly spin-labeled Tau187 + 90% unlabeled Tau187), such that statistically the closest distances are from the pair of spin-labels attached to the same Tau187 molecule. Effects from longer inter-molecular distances between spin-labels are corrected by dividing a background function from the time-domain DEER decay that is modulated by the weaker inter-molecular spin-spin interactions. Seeding assays conducted using spin-diluted Tau187 yielded similar ThT fluorescence intensity profiles as the aggregation assay using unlabeled Tau187 alone (Fig. S2). To prepare samples for DEER measurements, we induced tau aggregation by adding either heparin or 15% lysates for 24 hours, followed by dialysis (molecular weight cut off of 25 kDa) to remove excess monomer and reduce ^1^H concentration for DEER measurements. Using DEER, we derived the intra-tau spin-label distance distribution, P(*r*), from experimental data and reconstructed it with the DeerLab fitting algorithm.[48] Confidence intervals (95 we added heparin or 15% lysates, respectively, to induce aggregation for 24 h. Samples were then subjected to dialysis against a *D*_2_O buffer with 20 mM HEPES to remove excess monomer and to reduce ^1^H concentration to aid DEER measurements. We then derived the distribution of intra-tau spin-label distance, P(*r*), from the experimental time-domain DEER data, V(t), which is modulated by convolved dipolar coupling frequencies between pairs of spin-labels. The reconstruction of the distance distribution, P(*r*), from V(t) was determined using the recently published DeerLab fitting algorithm and software. The 95% confidence intervals were determined using the bootstrapping method with a sample size of 100.

To further assess the DEER data collected across samples, we utilized a brand-new data analysis methodology developed by Srivastava, Freed and coworkers, which uses a frequency pattern recognition technique that combines discretized continuous wavelet transform (CWT) with a structure similarity index measure (SSIM) analysis. This method relies on xxx By calculating the CWT, we can distinguish between the DEER signal, noise, and artifacts from the experiments by decomposing the DEER signal into different frequency components, thereby allowing a much more detailed comparison of the data than visually comparing the raw V(t) and the extracted P(*r*).[49] By utilizing the SSIM analysis, we can further compare the CWT of the DEER data because SSIM provides errors based on 1) perception as it accounts for the structural and textural information in addition to the magnitude of errors, and 2) saliency by giving more importance to the significant parts of the structures.[50] The result of this frequency pattern recognition is displayed as a similarity gradient plot, along with the SSIM index, to compare spectrograms of heparin-induced and CBD and PSP lysate-seeded fibrils (Fig. 5).

The DEER-derived distance distributions for three spin-label pairs on heparin-induced fibrils and the CBD and PSP lysate-seeded fibrils were compared with the simulated DEER distance distributions expected from the reported cryo-EM CBD and PSP folds (Fig. 4). The distance distributions between the spin-label pair across residues 351 & 373, referred to as P(*r*, 351-373), of the three different samples, are shown in Fig. 4a. The heparin fibrils (green) yielded the broadest distance distribution, which is distinct from the CBD (pink) and PSP (blue) lysate-seeded fibrils.

**Fig. 4.**
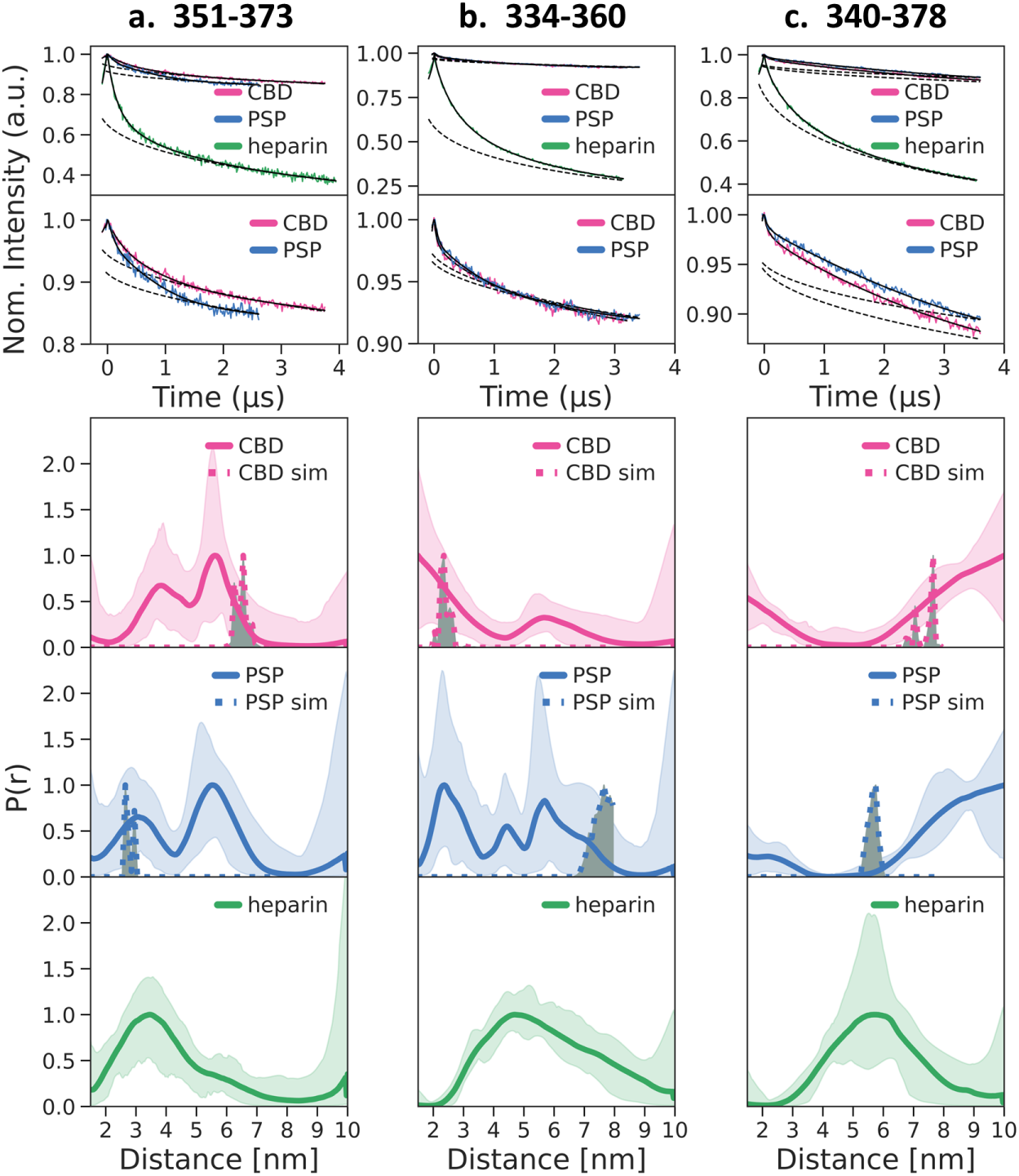
Intra-molecular DEER distance distribution, P(*r*), measured for heparin fibrils (green, solid), CBD lysate-seeded fibrils (pink, solid), and PSP lysate-seeded fibrils (blue, solid). Expected distances (dotted curves) were simulated from the published cryo-EM structures of the CBD fold (PDB: 6TJO; pink, dotted) and PSP fold (PDB: 7P65; blue, dotted) [19, 21] by the RotamerConvolveMD method [36] and are illustrated on the schematic view of these folds in Fig. 1. Three spin-label pairs were measured and simulated for (a) residues 351 & 373, (b) residues 334 & 360, and (c) residues 340 & 378. Replicates are shown in SI Figures, Fig. S5 to Fig. S7. The raw DEER signal V(t) are shown with the fit (in black) and the background fit (in black dotted).

The similarity gradient plot of the DEER signal from heparin-induced vs CBD lysate-seeded fibrils (SSIM 0.59), heparin-induced vs PSP lysate-seeded fibrils (SSIM 0.55), and PSP vs CBD lysate-seeded fibrils (SSIM 0.93) showed localized frequency differences with the SSIM indicating heparin-induced fibrils are vastly different from CBD and PSP lysate-seeded fibrils (Fig. 5a). Although the P(*r*) of the PSP and CBD lysate-seeded fibrils look nearly identical (SSIM 0.93), the similarity gradient plot shows differences in the higher frequency region (i.e., short distances region), validating the P(*r*) difference (see last figure of Fig. 5a). These results demonstrate that the heparin-induced fibrils are structurally unique and are more heterogeneous in the MTB region of the protein spanning residues 351-373. In contrast, the CBD and PSP lysate-seeded fibrils (in green and pink) exhibited a distinct shape for the distribution of the 351-373 distance, P(*r*, 351-373), featuring two dominant distance features with narrower widths compared to that of heparin fibrils, suggesting that multiple and distinct populations of ordered tau conformers are present.

**Fig. 5.**
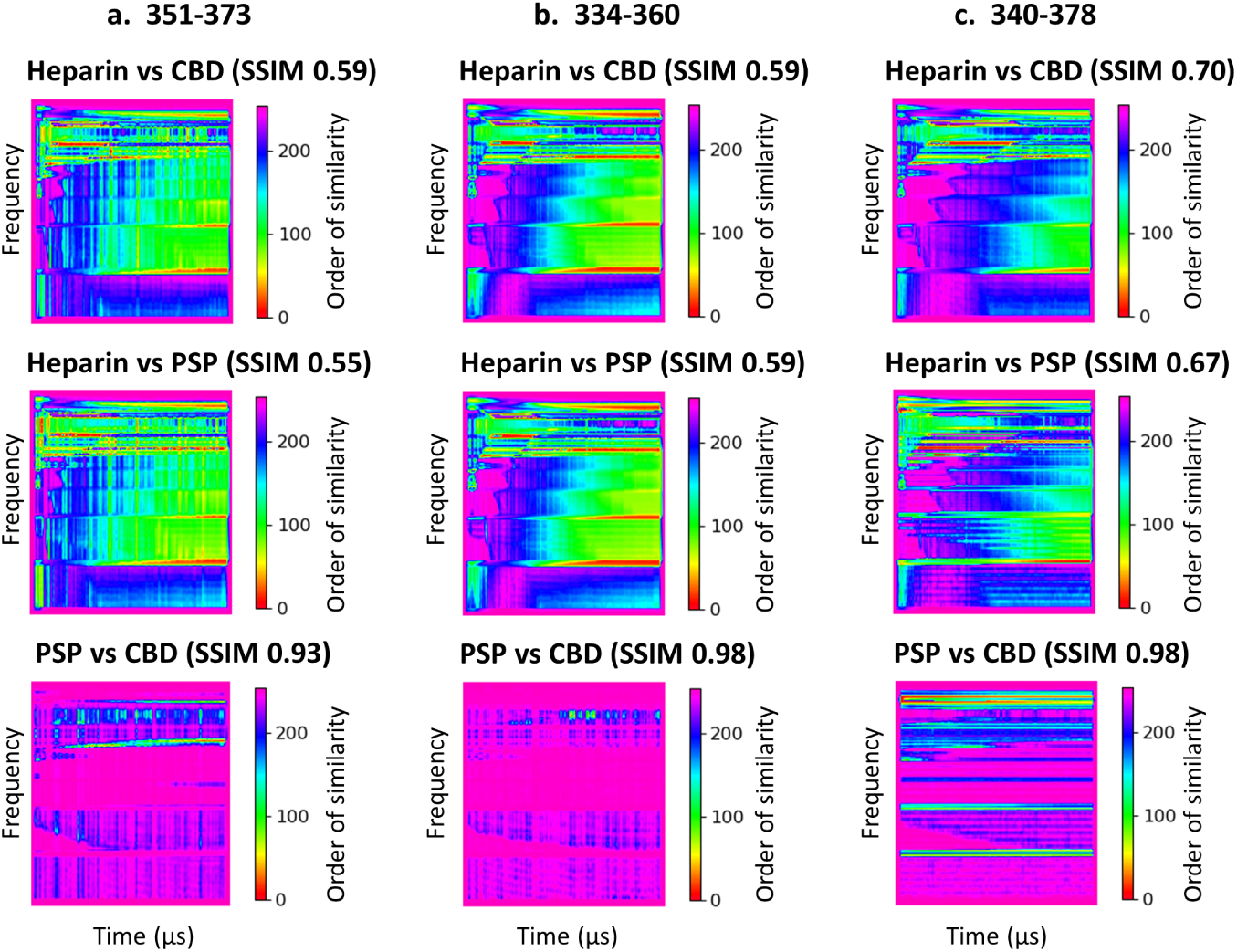
Comparison of the spectrograms of the CWT of raw DEER signal V(t) shown in Fig. 4. Similarity gradient plot of DEER signal of heparin vs CBD-seeded fibrils, heparin vs PSP-seeded fibrils, and PSP vs CBD-seeded fibrils labeled at (a) residues 351 & 373, (b) residues 334 & 360, and (c) residues 340 & 378. Spectrograms are shown in Fig. S8-S10. Spectrograms of replicates are shown in Fig. S11-S14. SSIM of replicates are shown in Table SI.

Despite the complexity of the observed P(*r*) shape, the patterns of the peaks representing the CBD vs PSP lysate-seeded fibrils are highly reproducible. Each P(*r*) was measured in triplicate (all data shown in Fig. S5 - 7) and their CWT was analyzed (all data shown in Fig. S8 - 14, Table S1), showcasing remarkable reproducibility between the repeats. The triplicates were derived from multiple cellular batches prepared at different times spanning multiple years, and the lysate-seeded samples were derived from at least three different *in vitro* seeding experiments performed on different batches of recombinant Tau187 using biologically different cell-passaged seeds. Critically, the shape for P(*r*) is clearly distinct between the CBD and PSP lysate-seeded fibrils. These results reveal that the cell-passaged seeds play an active role in templating and misfolding tau, with the two seeds inducing unique patterns for P(*r*).

An equally important observation is that each distinct pattern of P(*r*) is still broad, suggesting that the templating induced by seeding does not generate converged fibril structures with the commonly used seed material and seeding protocol by many researchers. Critically, a dominant peak near the predicted spin-label distance (indicated by dotted lines) suggests that a subpopulation of fibrils resembling the CBD and PSP disease folds may have formed. Specifically, a mean distance of 6.5 nm is expected for the CBD fold, while a dominant experimental peak is found at distances of 5-7 nm. In contrast, a mean distance of 3 nm is expected for the PSP fold, and again, one of the dominant experimental peaks is found spanning distances of 2-4 nm. Notably, this intensity at 2-4 nm is missing in the P(*r*, 351-373) for the CBD lysate-seeded tau fibril population.

We next analyzed the P(*r*, 334-360) data measured by DEER using Tau187 labeled across sites 344 & 360. Again, the heparin fibrils (green) yielded a distance distribution, which is distinct from the CBD (pink) and PSP (blue) lysate-seeded fibrils (Fig. 4b). This is further shown by the similarity gradient plot of DEER of heparin-induced vs CBD lysate-seeded fibrils (SSIM 0.59), heparin-induced vs PSP lysate-seeded fibrls (SSIM 0.59), and PSP vs CBD lysate-seeded fibrils (SSIM 0.98), showing localized frequency differences with SSIM indicating heparin-induced fibrils are vastly different from CBD and PSP lysate-seeded fibrils (Fig. 5b). The experimental P(*r*, 334-360) of CBD lysate-seeded fibrils (pink solid line) contains a peak comprising the expected distance distribution (pink dotted line) centered around 2.5 nm. The P(*r*, 334-360) of PSP lysate-seeded fibrils also has a dominant peak centered around 5-7 nm, which is closer to the expected distance of 7.5 nm. Although the P(*r*) of the PSP and CBD seeded fibrils look nearly identical (SSIM 0.98), the similarity gradient plot shows differences in the higher frequency region (i.e., short distances region), validating the P(*r*) difference where the CBD lysate-seed fibrils contain more high-frequency data than that of PSP lysate-seeded fibrils (see last figure of Fig. 5b). In contrast, the P(*r*, 334-360) of the heparin fibrils significantly differs from the measured P(*r*) of each of the two lysate-seeded fibrils, with no peaks present at distances above 6 nm. Again, the P(*r*) shape is remarkably reproducible across biological triplicates (in terms of the seeded tau fibril material; Fig. S6, S11-S14; Table S1), and is distinct between CBD and PSP lysate-seeded fibrils despite the greater complexity and width of the experimentally observed P(*r*) than commonly deemed interpretable.

The results for P(*r*, 340-378) also show a highly distinct P(*r*) pattern for the heparin-induced fibrils compared to the lysate-seeded tau fibril populations (Fig. 4c). This is supported by the similarity gradient plot of DEER signal of heparin-induced fibrils vs CBD lysate-seeded fibrils (SSIM 0.70), heparin-induced fibrils vs PSP lysate-seeded fibrils (SSIM 0.67), and PSP vs CBD lysate-seeded fibrils (SSIM 0.98) showing localized frequency differences with SSIM indicating heparin-induced fibrils are vastly different from CBD and PSP lysate-seeded fibrils (Fig. 5c). However this time, the P(*r*) between CBD or PSP lysate-seeded fibrils could not be distinguished. The major peak recorded from the CBD or PSP lysate-seeded fibrils is around 7-10 nm. This is slightly greater than the expected mean distance of 7.5 nm for the CBD fold, and much greater than the expected 5.5 nm for the PSP fold. Still, the distances contributing to the experimental P(*r*) data from CBD and PSP lysate-seeded fibrils were significantly greater than those found in heparin fibrils. Although the P(*r*) of the PSP and CBD seeded fibrils looks nearly identical (SSIM 0.98), the similarity gradient plot shows small differences across the frequency ranges (last figure in Fig. 5c). The measurements of P(*r*, 340-378) have been intrinsically more challenging compared to P(*r*, 351-373) and P(*r*, 334-360). This is because the expected distance range between sites 340 and 378 was the largest (5.5 to 8 nm) and is, therefore, close to the limitations of DEER with the given phase memory time, *T_m_*, of 4 *µ*s using Tau187 fibrils prepared under the conditions of this study. Additionally, site 378 lies at the most terminal position of the structure, with respect to the expected CBD or PSP fibril core [19, 21], compared to all other spin-labeled sites tested here. The Tau4RD expressed in the Tau4RD*LM-YFP cells also ends at residue 378, and the subsequent C-terminal residues present in both the CBD and PSP folds may be key to stabilizing the structures. It is, therefore plausible that the terminal segment around site 378 is not fully folded into the CBD or PSP cell-passaged seeds used here, resulting in additional broadening and uncertainty in the P(*r*) shape. Conventionally, the interpretation of P(*r*, 340-378) data given the very broad features spanning distances exceeding 7 nm as found with CBD and PSP lysate-seeded fibrils is not viable. However, the remarkable reproducibility of P(*r*) patterns between triplicates (Fig. S7, S11-S14; Table S1) demonstrates that the structural features, as seen through the dipolar coupling distributions between spin-labels at sites 340-378, are reproducible between the fibrils generated by each of the three methods.

When comparing the DEER-derived P(*r*) of Tau187 in the monomer and heparin-induced fibril state across sites 351-373, 334-360, and 340-378 (shown in Fig. S4), it is clear that the protein region spanning sites 340-378 is part of a semi-ordered fibril region and not simply an intrinsically disordered fuzzy coat in contrast to the P(*r*, 340-378) of heparin-induced fibril that exhibit a single peak. This result is not necessarily obvious, given that sites 334, 351, 360, 373, and 378 have not been reported to be part of any heparin-induced fibril fold according to cryo-EM [25], further highlighting the ability of DEER to report on less homogeneous and more dynamic protein folds within fibrils. Each P(*r*) of CBD or PSP lysate-seeded fibrils contain populations that feature greater distances between sites 351-373, 334-360, and 340-378, compared to heparin-induced fibrils, demonstrating that these sites are located at the outer core of a sub-population of tau fibrils and adopt more extended conformations, as expected from the PDB structure of the CBD and PSP folds. In other words, the *in vitro* CBD and PSP lysate-seeded fibrils contain sub-populations of fibrils that adopt structural features that are consistent with the tau structures found in CBD and PSP patients by cryo-EM. These findings indicate that cell-passaged CBD and PSP can transmit their structural properties to naïve tau by templated misfolding, at least in part, even though the biosensor cell lines used in this study express Tau4RD spanning residues 244-378, which is a few amino acids short of the tau segment that forms the PSP and CBD core. The results also show that seeding with cell-passaged patient-derived tau populates multiple fibril structures. It is also conceivable that the particle picking and class averaging approach of cryo-EM result in resolving only the dominant fibril sub-population, failing to capture other conformations that may exist in the human brain. These studies will benefit from experimental approaches to measure the complete ensemble of the fibril population that guide the discovery of mechanisms and conditions to achieve structural convergence to generate pathological tau fibrils.

## 3 Discussion

In this study, we successfully used cell-passaged CBD and PSP patient samples to induce recombinant tau fibrillization without the addition of any co-factors or the use of sonication. Moreover, the resulting fibrils were able to reinfect the Tau4RD*LM-YFP cells showing isoform specificity (Fig. 3c & d). Using DEER-based distance measurements at sufficiently high-resolution and sub-nanometer levels, we found that the CBD and PSP lysate-seeded fibrils are structurally distinct from one another, as well as from heparin-induced tau fibrils (Fig. 4). Together, these results indicate that we can propagate and measure distinct fibril structures, but that additional factors must be controlled to achieve structural convergence. Our study shows that DEER is an effective adjunctive tool to cryo-EM to evaluate strain-specific tau misfolding for any experimental condition, independent of whether convergence of fibril structure is achieved or not.

Our results are consistent with the predominant hypothesis that tau fibril propagation uses the prion mechanism to self-template distinct conformations or strains. While cryo-EM has revealed a diverse array of tau fibril conformations [19–23, 23, 44], the ability to replicate these structures upon seeding has only recently been completed at high-resolution. An exciting study was published recently by Tarutani *et al.* reporting the first cryo-EM structures of cell-passaged insoluble tau fibrils [51]. Complementing this exciting work, we show here that DEER can uniquely capture an ensemble of structures within a larger population, not just a homogeneous or dominant sub-population.

Previously, morphological and biochemical properties were used to demonstrate that distinct tau conformations exist [13, 14, 27, 27, 28, 33], and similar features between the seeded fibrils and the starting seed, or tau prion, have been used to determine the fidelity of tau templating [13, 14, 27–33, 35].

For example, Kaufman *et al.* [10] isolated and characterized 18 tau strains that originated from either recombinant protein, tauopathy mouse brain, or deceased human tauopathy patient samples. The isolated strains were categorized using multiple approaches, including inclusion morphology, detergent solubility, seeding activity, proteolytic digestion, and toxicity. After inoculating transgenic mouse brains with the 18 strains, the unique patterns of neuropathological lesions induced by each strain were faithfully transmitted, indicating that the seeded fibrils in mice are similar to the starting material in the inocula. In a more recent study, Xu *et al.* [33] showed that AD patient samples may be used to template recombinant tau misfolding, resulting in fibrils that adopt the same conformation. While their analyses lacked confirmation via cryo-EM, the conformation-dependent antibodies DMR7 and MC1 were able to bind to the AD-induced fibrils. In contrast, others have found that the AD seeded fibrils adopt conformations that are distinct from the starting material, as determined by unique protease digestion patterns and macroscopic morphology of induced fibrils [13, 52]. The current state of the literature shows that high-resolution tools are needed to effectively guide this debate regarding strain-specific templated misfolding of tau. Studies such as ours will contribute to the critical ongoing debate on the molecular basis of tau prions and shape-selected fibril formation.

Using DEER and frequency pattern recognition, we detected partial distance features of seeded tau fibrils that are similar to the expected distances measured from the cryo-EM structures of CBD and PSP tau. These findings indicate that we achieved some convergence with these structures using seeding, even when using imperfect cellular seed material. One might view these results only as partial success given that the fibril structures did not converge to a single structure resembling the CBD or PSP fold. While achieving complete convergence to disease fold is an ultimate goal, the data presented here highlight a critical and unique strength of DEER, that it is capable of capturing the entire population of tau folds within the formed fibrils. No other experimental technique could reveal that tauopathy seeds derived from cell lysates do, on the one hand, generate the originally targeted disease fold, while on the other hand also report that many other fibril structures coexist with the disease fold. The ability to capture the entire conformational population is powerful because this tool offers the opportunity to learn what dominant and relevant factors are needed to achieve conformation selection and convergence. Importantly, there are a few factors that may account for this partial convergence, rather than complete replication of fibril structure. First, the cell-passaged seeds were generated in cells expressing Tau4RD which lacks the N- and C-termini. Although the Tau4RD in cells may be templated by human materials, residues outside of the MTBR (such as the rest of C-terminal region after residue 378) may contribute to tau folding, strain propagation, and/or structural stabilization. As a result, the CBD and PSP folds may only partially form in cells. This is potentially reflected in the indistinguishable P(*r*, 340-378) data measured from the CBD and PSP lysate-seeded fibrils. Moreover, despite the use of an 18-amino acid flexible linker, the fusion of Tau4RD to a fluorescent protein may induce steric hindrance that inhibits or alters tau aggregation, as reported recently [53]. However, Tau4RD is still important because it is so far the most commonly used construct in cellular seeding assays [11, 24, 54–64]. We, therefore, need to know how to interpret the wealth of critical data previously reported, no matter what the ultimate verdict is (e.g., whether the 4RD seeds do or do not faithfully template disease folds). Furthermore, truncated tau remains biologically relevant because fragments of tau have been identified in AD patient brain samples [63], and in patient cerebrospinal fluid [65–67]. Moreover, the linker sequence connecting tau and GFP in the studies by Kaniyappan *et al* (GAPGSAGSAAGSG) is not the same linker sequence used in the cell model used here (EFCSRRYRGPGIHRSPTA). It is unclear how differences in the length or flexibility of these two linker sequences may contribute to our findings, and therefore how translatable the work by Kaniyappan *et al* is to the model system used here. In light of this discrepancy in the literature, others have shown that the presence of a GFP tag on *α*-synuclein does not interfere with *in vitro* fibrillization [68]. Therefore, it remains an open question whether the cell-passaged seeds are disrupted by the YFP tag and the protein truncation.

Second, the cell-free seeding conditions used in this study may not favor the propagation of specific disease-relevant strains, resulting in the formation of multiple conformers. For example, no disease-relevant PTMs are present on the recombinant tau used here. Additionally, the exact makeup of the solution conditions, including molecular chaperones, crowding agents, co-factors, and even salt type and concentration likely play major roles in dictating the tau folding pathway. In fact, a recent study using cryo-EM showed that simply changing salt types and salt concentration in the seeding buffer dramatically alters the shape of the resulting tau fibrils [45]. By systematically and empirically changing the buffer conditions, the authors successfully generated recombinant tau fibrils identical to the conformations reported for AD and CTE tau. Therefore, further optimization of the seeding conditions may be needed, however, it is clear that our approach offers the opportunity to understand the underlying mechanism(s) for shape selection in tau seeding.

We need to consider two important factors when interpreting cryo-EM structures of tau fibrils. The cryo-EM structures are generated using samples from deceased patients and therefore represent the end stage of the disease. It is possible that the process of tau misfolding in a living patient is more dynamic. Hence, the heterogeneous nature of the tau fibril population reflected in our dataset captured from fibril samples 24 h after seeding may have more biological relevance for earlier stages of tau misfolding. Additionally, the class averaging required to resolve these structures likely selects for the most abundant conformations among a population of fibrils. However, as has been shown for the prion protein, there may be a mixture, or cloud, of conformations present in patients with CBD and PSP. If this is the case, it is less likely that cryo-EM studies will resolve the less abundant tau species, which we may be detectable by DEER, demonstrating the need for complementary structural biology tools.

Finally, an important limitation to our current work is the use of one CBD and one PSP patient sample to generate Tau4RD*LM-YFP cells that stably propagate tau aggregates. While this approach was taken to support the initial establishment of the methodology and approach reported here, it is possible that by using a single patient sample, we are unable to fully capture the diversity in P(r) that exists across tau misfolded for the two diseases. Future studies will focus on understanding the patient-to-patient variability in tau misfolding that may exist, which will be done using cell lines that avert some of the caveats discussed above.

## 4 Conclusion

We demonstrate that the recombinant tau fibrils induced by CBD and PSP cell-passaged seeds are structurally distinct from heparin-induced fibrils and that a sub-population of the lysate-seeded fibrils resemble the CBD and PSP fibrils isolated from human patient samples. These findings validate the application of DEER to investigate the self-templating process during tau propagation at high-resolution. Moreover, these data will also be important for guiding the development of strain-specific positron emission tomography (PET) ligands [69]. Given the conformational diversity of tau fibrils across tauopathy patients, it is unsurprising that previously developed PET probes bind some conformations with a much higher affinity than others (such as the APN-1607 binding to tau inclusions from AD and PSP patients)[70, 71]. In order to improve the diagnosis of tauopathies in living patients, additional PET ligands specific to each conformation are needed.

## 5 Methods

Parts of the experimental procedures outlined in this section were previously described in the doctoral thesis of the first author, Zhikai Zeng, titled “Seeking the Structural Basis of Tau Seeding”[72].

### 5.1 Human patient samples

The CBD patient sample used was provided by the NIH NeuroBioBank. This sample was from a 65-year-old male patient. The PSP patient sample used was provided by the Massachusetts Alzheimer’s Disease Research Center. This sample was from a 70-year-old female patient. Fresh-frozen human tissue was used to create a 10% (wt/vol) homogenate using calcium- and magnesium-free 1X Dulbecco’s phosphate-buffered saline (DPBS) using an Omni Tissue Homogenizer (Omni International).

### 5.2 Cell line development

The human tau DNA sequence encoding residues 244-378 with the P301L and V337M mutations (based on the longest tau isoform, 2N4R) fused to enhanced yellow fluorescent protein by an 18 amino acid flexible linker (EFC-SRRYRGPGIHRSPTA) was synthesized and cloned into the pcDNA3.1(+) expression vector by GenScript. The Tau4RD*LM-YFP sequence was then subcloned into the pIRESpuro3 vector (Takara) using EcoRV (5’) and NotI (3’). The same protocol was used to construct a plasmid containing the repeat domain of 3R tau (residues 244-274,306-378) containing the L226V and V337M mutations. Gene sequence and insertion were confirmed by Sanger sequencing before subsequent use.

HEK293T cells (ATCC) were cultured in Dulbecco’s modified Eagle’s medium (DMEM; Corning) supplemented with 10% fetal bovine serum (FBS), 100 *µ*g/mL penicillin, and 100 *µ*g/mL streptomycin (ThermoFisher), referred to hereafter as complete media. Cultures were maintained in a humidified atmosphere of 5% CO2 at 37 °C. Cells were plated at a density of 5.7 X 10^5^ cells per well in a 6-well plate overnight in complete media before adding 1 *µ*g of plasmid DNA incubated with 3.5 *µ*L Lipofectamine 2000 (ThermoFisher) for 20 min. Stable cells were selected in complete media containing 1 *µ*g/mL puromyocin (ThermoFisher) for 48 h before generating monoclonal lines by limiting dilution of polyclonal cells in 384-well plates. The resulting monoclonal lines were frozen in liquid nitrogen. Lysates from the lines were collected in 1X radioimmunoprecipitation assay (RIPA) buffer containing 50 mM Tris-HCl, pH 7.5 (ThermoFisher), 150 mM NaCl (Sigma), 5 mM EDTA (ThermoFisher), 1% nonidet P-40 (ThermoFisher), 0.5% deoxycholate (ThermoFisher), and 0.1% sodium dodecyl sulfate (SDS; ThermoFisher). Cell lysates in RIPA buffer were frozen, thawed, and clarified using two low-speed spins (500 x g for 5 min followed by 1,000 x g for 5 min). Total protein was measured in the supernatants via bicinchoninic acid (BCA) assay (Pierce). To compare the expression of tau-YFP across the clones, a total of 10 *µ*g total protein was prepared in 1X NuPAGE LDS loading buffer and boiled for 10 min. Samples were loaded onto a 10% Bis-Tris gel (ThermoFisher) and SDS-PAGE was performed using the MES buffer system. Protein was transferred to a polyvinylidene fluoride (PVDF) membrane. The membrane was blocked in blocking buffer [5% (wt/vol) nonfat milk in 1X Tris-buffered saline containing 0.05% (vol/vol) Tween 20 (TBST)] for 30 min at room temperature. Blots were incubated with primary antibody for GFP (1:10,000; Abcam) in block buffer overnight at 4 °C. Membranes were washed three times with 1X TBST before incubating with horseradish peroxidase-conjugated goat antib-rabbit secondary antibody (1:10,000; Abcam) diluted in blocking buffer for 1 h at 4 °C. After washing blots three times in 1X TBST, membranes were developed using the enhanced chemiluminescent detection system (Pierce) for exposure to X-ray film. This same protocol was used to evaluate tau-YFP expression in uninfected vs infected cells.

To generate cells that stably propagate tau prions, fresh frozen human brain tissue samples from one CBD and one PSP patient sample were used. Tau prions were isolated from the samples using sodium phosphotungstic acid (NaPTA; Sigma) as described [11]. Isolated protein pellets were stored at 4 °C. Tau4RD*LM-YFP cells were infected with tau prions isolated from the CBD or PSP patient samples as described below. Monoclonal subclones stably propagating tau prions were generated by isolated Tau4RD*LM-YFP cells by limiting dilution of a polyclonal cell population in 384-well plates. Clones were selected by the presence of YFP-positive aggregates in >95% of cells in the culture. Resulting monoclonal lines were frozen in liquid nitrogen.

### 5.3 Cellular tau prion bioassay

Tau4RD*LM-YFP cells were plated in 384-well plates with black polystyrene walls (Greiner) with 0.012 *µ*g Hoechst 33342 (ThermoFisher) at a density of 4,000 cells/well. Tau3RD*VM-YFP cells were plated using the same conditions at a density of 2,750 cells/well. Cells were incubated at 37 °C for 2 to 4 h to allow adherence to the plate. To infect Tau4RD*LM-YFP cells, PTA-precipitated patient samples diluted 1:5 in 1X DPBS or 0.05 *µ*M fibrils in 1X DPBS were incubated with Lipofectamine 2000 2% final volume; ThermoFisher) for 1.5 h at room temperature. OptiMEM (78% final volume; ThermoFisher) was added before each sample was plated in six replicate wells. To infect Tau3RD*LM-YFP cells, PTA-precipitated patient samples diluted 1:20 in 1X DPBS or 0.05 *µ*M fibrils in 1X DPBS were incubated with Lipofectamine 2000 1% final volume; ThermoFisher) for 1.5 h at room temperature. OptiMEM (79% final volume; ThermoFisher) was added before each sample was plated in six replicate wells. Plates were then incubated at 37 °C in a humidified atmosphere of 5% (vol/vol) CO_2_ for 4 days before imaging on the Molecular Devices XLS.Images of both the DAPI and YFP channels were collected from five different regions in each well. The images were analyzed using an algorithm built in Custom Module Editor (Molecular Devices) to identify intracellular aggregates only in living cells. Data reported as fluorescence/cell X 10^6^ arbitrary units (A.U.).

### 5.4 Tau187 and 0N4R expression and purification

N-terminal truncated, microtubule-binding domain containing Tau187 (residues 255-441 with a His-tag at the N-terminus) and 0N4R were used for *in vitro* studies. Mutated variants of Tau187 were prepared using site-direct mutagenesis: cysteine-less (cysless) Tau187 P301L/Q351C/T373C contains C291S, C322S, Q351C and T373C; cysless Tau187 P301L/G334C/I360C contains C291S, C322S, G334C and I360C; cysless Tau187 P301L/K340/F378C contains C291S, C322S, K340 and F378C. Cysless Tau187 P301L and cysless 0N4R P301L constructs were made with C291S and C322S mutations.

The cloning, expression, and purification of Tau187 have been previously described [73]. Shortly, the gene encoding desired Tau187 tau or 0N4R tau was cloned into a pET-28a vector and was tranfected into E. coli. BL21 (DE3) competent cells (Novagen). E. coli BL21 (DE3) cells were cultured from frozen glycerol stock or from plates overnight in 10 mL luria broth (LB) which was used to inoculate 1 L of fresh LB. Culturing and inoculation were performed at 37 °C with shaking of 200 rpm. At OD 600 of 0.6-0.8, Tau187 variant expression was induced by incubation with 1 mM isopropylß-D-thiogalactoside (Sigma Aldrich) for 2-3 h. Cells were harvested by centrifugation for 10 min at 5000 × g (Beckman J-10; Beckman Instruments,Inc.), and the pellets were stored at −20 °C until further use. Cell pellets were resuspended in lysis buffer (Tris-HCl pH 7.4, 100 mM NaCl, 0.5 mM DTT, 0.1 mM EDTA, 1mM PMSF) with 1 Pierce protease inhibitor tablet (Thermo Fisher). Lysis was initiated by the addition of lysozyme (2 mg/ml), DNase (20 *µ*g/ml), and MgCl_2_ (10 mM) and incubated for 30 min on ice. Lysate was then heated to 65 °C for 13 min, cooled on ice for 20 min and then centrifuged to remove the precipitant. The supernatant was loaded onto a Ni-NTA agarose column pre-equilibrated with wash buffer A (20 mM sodium phosphate pH 7.0, 500 mM NaCl, 10 mM imidazole, 100 *µ*M EDTA). The column was then washed with 20 ml of buffer A, 15 ml buffer B (20 mM sodium phosphate pH 7.0, 1 M NaCl, 20 mM imidazole, 0.5 mM DTT, 100 *µ*M EDTA). Purified Tau187 was eluted with buffer C (20 mM sodium phosphate pH 7.0, 0.5 mM DTT, 100 mM NaCl) supplemented with varying amounts of imidazole increasing from 100 mM to 300 mM. The protein was then concentrated via Amicon Ultra-15 centrifugal filters (MWCO 10 kDa; Millipore Sigma) and the buffer was exchanged into final buffer (20 mM ammonium acetate buffer at pH 7.0, or 20mM HEPES at pH 7.4) by PD-10 desalting column (GE Healthcare). To further purify the protein, size exclusion chromatography was conducted by injecting 2-5 mL sample onto HiLoad 16/600 Superdex 200 Column (GE Healthcare Life Sciences) connected to a BioRad NGC Quest 10 FPLC system, and 1.5 mL fractions were collected with a BioFrac fraction collector (BioRad). Prior to sample injection, the column was washed by 130 mL degassed Milli-Q water at a flow rate of 0.8 mL/min and equilibrated with 160 mL degassed working buffer (20 mM ammonium acetate buffer at pH 7.0, or 20mM HEPES at pH 7.4) at 0.8 mL/min. Samples were eluted from the column with 150 mL working buffer at 0.6 mL/min after sample injection. Fractions of each elution peaks were collected right after elution. Elution peak assignment was done by comparing with the elution profile of purified Tau187 monomers. Fractions corresponding to monomer were concentrated using Amicon Ultra-4 centrifugal filters (MWCO 10 kDa; Millipore Sigma). The final protein concentration of Tau187 was determined by UV-Vis absorption at 274 nm using an extinction coefficient of 2.8 *cm^−^*^1^*mM ^−^*^1^ calculated from absorption of Tyrosine. The final protein concentration of 0N4R was determined by UV-Vis absorption at 274 nm using an extinction coefficient of 7.4 *cm^−^*^1^*mM ^−^*^1^ calculated from absorption of Tyrosine.

### 5.5 Protein spin-labeling

Protein was spin-labeled using MTSL ((1-Acetoxy-2,2,5,5-tetramethyl-*δ*-3-pyrroline-3-methyl) Methanethiosulfonate) purchased from Toronto Research Chemicals. Prior to labeling, samples were treated with 5 mM DTT, which was removed using a PD-10 desalting column. Then, 10× to 15× molar excess MTSL to free cysteine was incubated with the protein at 4 °C overnight. Excess MTSL was removed using a PD-10 desalting column. Labeling efficiency, defined as the molar ratio of tethered spin-labels over the cysteines, was measured to be 50-60% for double-cysteine mutants.

### 5.6 Heparin-induced tau aggregation

Cell-free tau aggregation was carried out in the working buffer (20mM HEPES at pH 7.4) at 37 °C. 20 *µ*M ThT dye was added into 50 *µ*M Tau187 in Corning™ 384-Well Solid Black Polystyrene Microplates (Thermo Fisher Scientific). Then, heparin was mixed in the samples at the mole ratio of 4:1 (Tau187:heparin) to induce aggregation. ThT fluorescence was monitored by Bio-Tek Synergy 2 microplate reader (excitation 440/30, emission 485/20, number of flash 16, gain 50) over a period of 24h at 37 °C. One data point was taken every 3 minutes. Measurements were done in triplicate. The figures showed the average and standard deviations.

### 5.7 Cell-free seeding assay

5% to 15% (protein mass percentage) of CBD and PSP cell-passaged seeds were incubated with 50 *µ*M cysless P301 Tau187 in 20 mM HEPES buffer to make seeded fibrils. 20 *µ*M ThT was added to the mixture and the fluorescence was monitored by Bio-Tek Synergy 2 microplate reader using the same setting and as heparin-induced aggregation. One data point was taken every 3 minutes. Measurements were done in triplicate. The figures showed the average and standard deviations.

### 5.8 TEM

TEM grids were plasma cleaned (for 20s at 60W power) with the shiny side up and used within 1 hour for preparation. 1 drop (5ul) of fibril sample and two drops (5ul each) of 2 percent UA (Uranyl Acetate) were placed on a parafilm for each grid. The shiny side was then floated on the fibril sample for 1 minute and was blotted once with Wattman filter paper. The grid was then touched with one drop of UA and blotted immediately. It was then placed on the second UA drop for 1 minute and then blotted well. The grids were dried for 2-3 hours, before imaging. Images were collected on a Talos F200X (Thermo Fisher Scientific) operating at 200 kV and equipped with a Ceta II CMOS 4k x 4k camera for high-quality HRTEM imaging.

### 5.9 Double Electron Electron Resonance (DEER)

Double-cysteine Tau187 was expressed and spin-labeled as doubly-labeled Tau187 that contains two spin-labels in order to probe distances between two target residues (residues 351 and 373, residues 334 and 360, and residues 340 and 378). Cysless Tau187 was expressed and purified to avoid disulfide bonding. doubly-labeled Tau187 and cysless Tau187 were stored in 20 mM HEPES in H_2_O. A 1:10 molar ratio doubly-labeled Tau187:cysless Tau187 sample of 57 *µ*M doubly-labeled Tau187 and 570 *µ*M cysless Tu187 was incubated with 157 *µ*M heparin at 37 °C 24 h to prepare heparin fibrils. As for CBD and PSP lysate-seeded fibrils, a 1:10 molar ratio doubly-labeled Tau187:cysless Tau187 sample of 200 *µ*M doubly-labeled Tau187 and 2000 *µ*M cysless was incubated with 15% (mass) CBD or PSP cell-passaged seeds at 37 °C for 24 h to prepare seeded fibrils. 50 *µ*L of fibrils were then dialyzed to D_2_O-based buffer ((20mM HEPES at pH 7.4) at room temperature for 6 h using Pur-A-Lyzer™ Mini Dialysis Kit (Mini 25000, MWCO 25 kDa). 35 *µ*L samples were then mixed with 15 *µ*L D8-glycerol (30% volume) before transferring to a quartz tube (3 mm o.d., 2 mm i.d.) and frozen using liquid nitrogen.

The DEER experiments were performed with a pulsed Q-band Bruker E580 Elexsys spectrometer, equipped with a Bruker QT-II resonator and a 300 W TWT amplifier with an output power of 20 mW for the recorded data (Applied Systems Engineering, Model 177Ka). The temperature of the cavity was maintained at 65 K using a Bruker/ColdEdge FlexLine Cryostat (Model ER 4118HV-CF100). The bridge is equipped with an Arbitrary Wave Generator to create shaped pulses for increased sensitivity. The following DEER pulse sequence was used: *π_obs_*/2 – *τ*_1_ – *π_obs_* – (*t* - *π_pump_*) – (*τ*_2_ - *t*) – *π_obs_* – *τ*_2_ – echo. Experiment dipolar signal, V(t), was recorded as the integral of the refoucsed echo as a function of time delay, *t*. Rectangular observe pulses were used with lengths set to *π_obs_*/2 = 10–12 ns and *π_obs_* = 20–24 ns. A chirp *π* pump pulse was applied with a length of 100 ns and a frequency width of 60 MHz. The observed frequency was 90 MHz higher than the center of the pump frequency range. *τ*_1_ was set to 180 ns for heparin samples and set to the first harmonic in the two-pulse deuterium ESEEM trace (126-128 ns) for cell seeded samples. *τ*_2_ was set according to the SNR profile of the dipolar signal. The data was acquired with Δ*t* of 16 ns, 16-step phase cycling, and signal averaged until desirable SNR was obtained.

### 5.10 DEER data analysis

All DEER time traces were transformed into distance distributions using DeerLab software package for Python. The time traces were phase corrected and truncated by 300 ns to remove possible "2+1"-artifact. One-step analysis was done using the DeerLab fit function with the following models: ex-4deer model with t1, t2, and pulselength set to experiment parameters, bg-strexp model with the stretch parameter freezed to 1.5 (base on previous measurements of dimensions of singularly labeled samples), and dipolarmodel using Tikhonove regularization. The uncertainty analysis was done using bootstrapping method with 100 samples. The time domain fitting results are presented with the fitting to the primary data and the background fit, and the distance distribution fits are presented with 95% confidence intervals.

### 5.11 DEER Frequency Pattern Recognition

All DEER time traces were decomposed into frequency components using discretized continuous wavelet transform (CWT). The calculated CWT was then converted into spectrograms measuring the range of frequencies present along the DEER trace. These spectrograms were compared using structure similarity index measure (SSIM) analysis by 1) numerical SSIM index, where SSIM index < 1 denotes a meaningful difference in frequency profile and by 2) similarity gradient plots, where the location with frequency differences can be visually identified.

### 5.12 Statistical analysis

Cell infection data are presented as mean *±* SD. The value represents the averages of five images collected from each well of a 384-well plate. Technical replicates for each sample were averaged across six wells. Statistical comparisons between control and diseased patient samples, and between monomer and fibrils, were performed using a one-way ANOVA with a Dunnett post-hoc analysis. Statistical significance for all tests was determined with a *P* value < 0.05.

## 6 Supplementary information

SI is attached as PDF.

## 7 Acknowledgments

The authors thank the DeerLab developers [48] for making an enhanced DEER analysis software available and Dr. Madhur Srivastava for continued discussion and support for DEER data analysis. Human tissue was obtained from the NIH NeuroBioBank and the Massachusetts Alzheimer’s Disease Research Center.

## 9 Author contributions

Z Zeng carried out the measurements and led the paper writing K. Tsay carried out the DEER measurements and helped lead the paper writing V. Vijayan carried out measurements and assisted in paper writing M. Frost did the seeding measurements S. Prakash did the DEER data analysis A. Quddus supported with measurements A. Albert supported with measurements M. Vigers supported intellectually and with assistance to measurements M. Srivastava guided and developed the DEER data analysis tool A. Woerman guided the cell seeding assay and helped write the paper S. Han guided the entire study, led the collaboration and helped write the paper

## 10 Data availability

The authors declare that the data supporting the findings of this study are available within the paper and its Supplementary Information files. Should any raw data files be needed in another format they are readily available from the corresponding authors upon reasonable request.

Editorial Policies for:

Springer journals and proceedings: https://www.springer.com/gp/editorial-policies

Nature Portfolio journals:

https://www.nature.com/nature-research/editorial-policies

*Scientific Reports*:

https://www.nature.com/srep/journal-policies/editorial-policies

BMC journals: https://www.biomedcentral.com/getpublished/editorial-policies

