## Supplemental Information for "Passaging Human Tauopathy Patient Samples in Cells Generates Heterogeneous Fibrils with a Subpopulation Adopting Disease Folds"

### Table of contents


| SI No. | Contents | Page No. |
|---|---|---|
| 1 | **Figure S1.** Uncropped full gel for Figure 2C. | 2 |
| 2 | **Figure S2.** ThT fluorescence of (a) 0N4R tau fibrillization. (b) recombinant Tau187 fibrillization induced by cell-passaged CBD seed. | 3 |
| 3 | **Figure S3.** Negative-stain TEM of seeded fibrils. | 4 |
| 4 | **Figure S4.** P(r) comparison between monomer and heparin induced fibrils. | 5 |
| 5 | **Figure S5.** Triplicates of P(r) measurements for residues 351 & 373 | 6 |
| 6 | **Figure S6.** Triplicates of P(r) measurements for residues 334 & 360. | 7 |
| 7 | **Figure S7.** Triplicates of P(r) measurements for residues 340 & 378. | 8 |
| 8 | **Figure S8.** Pattern Analysis of P(r) measurements for residues 351 & 373. | 9 |
| 9 | **Figure S9.** Pattern Analysis of P(r) measurements for residues 334 & 360. | 10 |
| 10 | **Figure S10.** Pattern Analysis of P(r) measurements for residues 340 & 378. | 11 |
| 11 | **Figure S11.** Spectrograms from CWT calculation of monomers. | 12 |
| 12 | **Figure S12.** Spectrograms from CWT calculation of heparin induced fibrils. | 13 |
| 13 | **Figure S13.** Spectrograms from CWT calculation of cell-passaged CBD seeded fibrils. | 14 |
| 14 | **Figure S14.** Spectrograms from CWT calculation of cell-passaged PSP seeded fibrils. | 15 |
| 15 | **Table S1.** SSIM index of each spectrogram comparison to one set control spectrogram. | 16 |

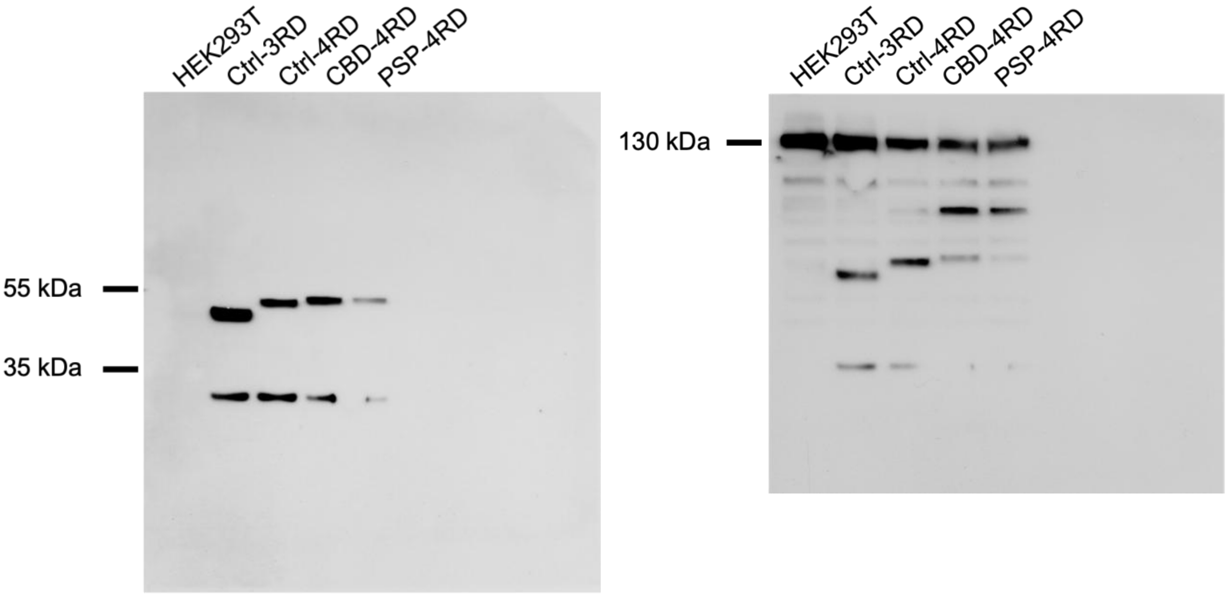


**Figure S1.** Uncropped blots of lysates collected in RIPA buffer from naïve HEK293T cells, Tau3RD*VM-YFP cells, Tau4RD*LM-YFP cells, and Tau4RD*LM-YFP cells stably infected with CBD or PSP tau prions shown in Fig. 2c. Left, membrane probed for the presence of the tau-YFP fusion protein using the GFP primary antibody. Right, membrane probed for vinculin as a loading control.

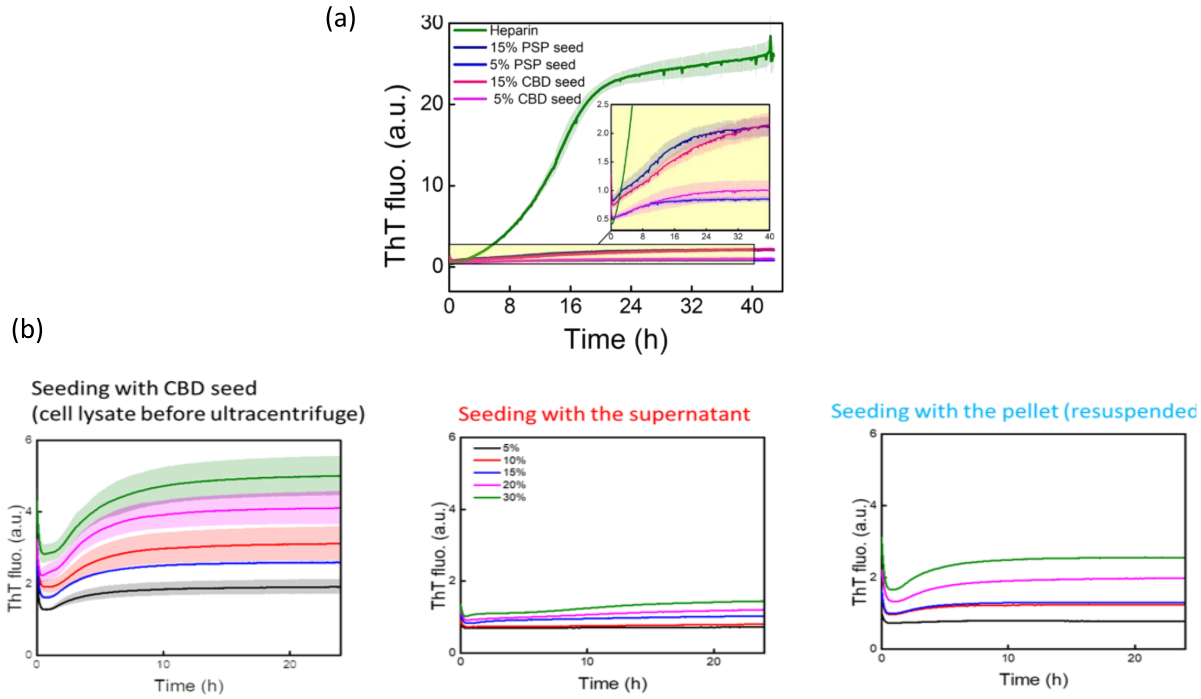


**Figure S2.** (a) ThT fluorescence of tau fibrillization induced by heparin (green), or cell-passaged seeds (lysates) from CBD (pink) or PSP (blue) patient samples for 0N4R construct. The lysate-seeded aggregation generated a relatively low maximum ThT; an enlarged curve is shown in the inset. 0N4R (25 µM) was mixed with stoichiometric amounts of heparin (8.25 µM, 17% by mass) or cell-passaged lysates [either 5% (lighter-color lines) or 15% by mass (darker-colored lines)] and was aggregated in the presence of ThT at 37°C. (b) ThT fluorescence of recombinant Tau187 fibrillization induced by cell-passaged CBD seed. Tau187 fibrillization in the presence of ThT at 37°C using CBD-seeded lysate (left) without treatment (middle) supernatant following ultracentrifugation, and (right) resuspended pellet following ultracentrifugation.

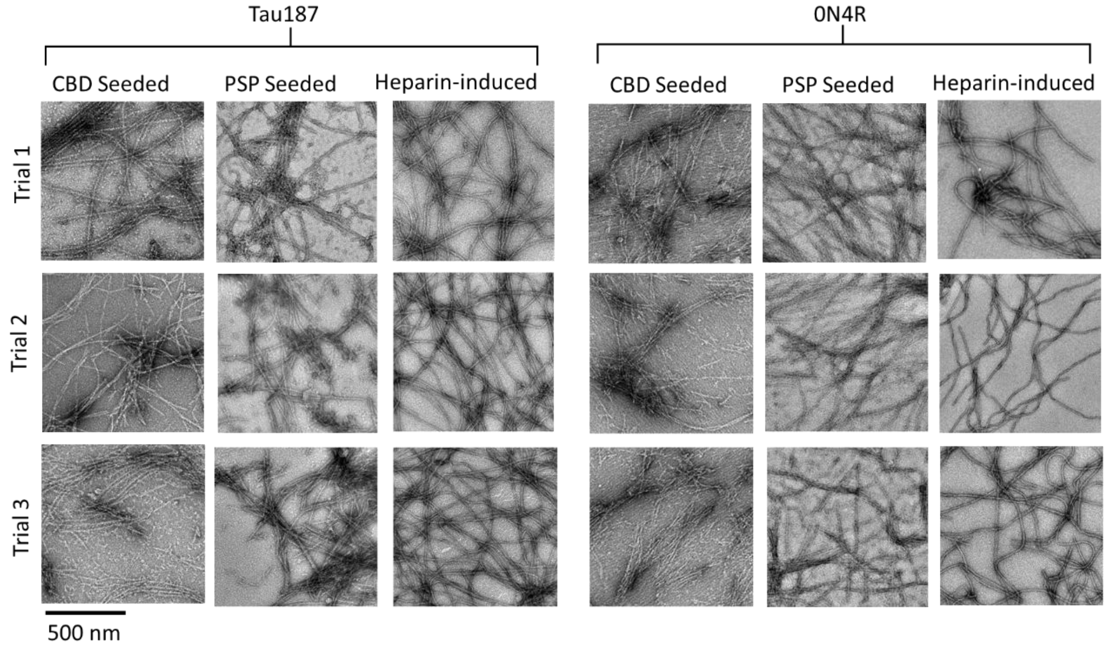


**Figure S3.** Negative-stain TEM of seeded fibrils. Triplicates (Trials 1-3) of negative-stain TEM images are shown for Tau187 (left) and 0N4R (right) seeded fibrils. Fibrillization was induced by CBD lysate (left), PSP lysate (middle), or heparin (right).

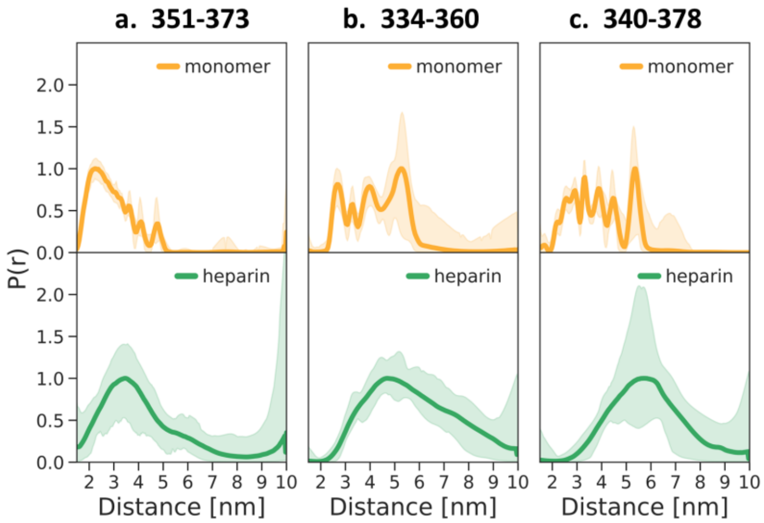


**Figure S4.** P(r) comparison between monomer and heparin induced fibrils. Measurements from residues (a) 351-373, (b) 334-360, and (c) 340-378. Technical replicates of P(r) measurements for residues 351 & 373 are in **Fig. S5**, residues 334 & 360 are in **Fig. S6**, and residues 340 & 378 are in **Fig. S7**.

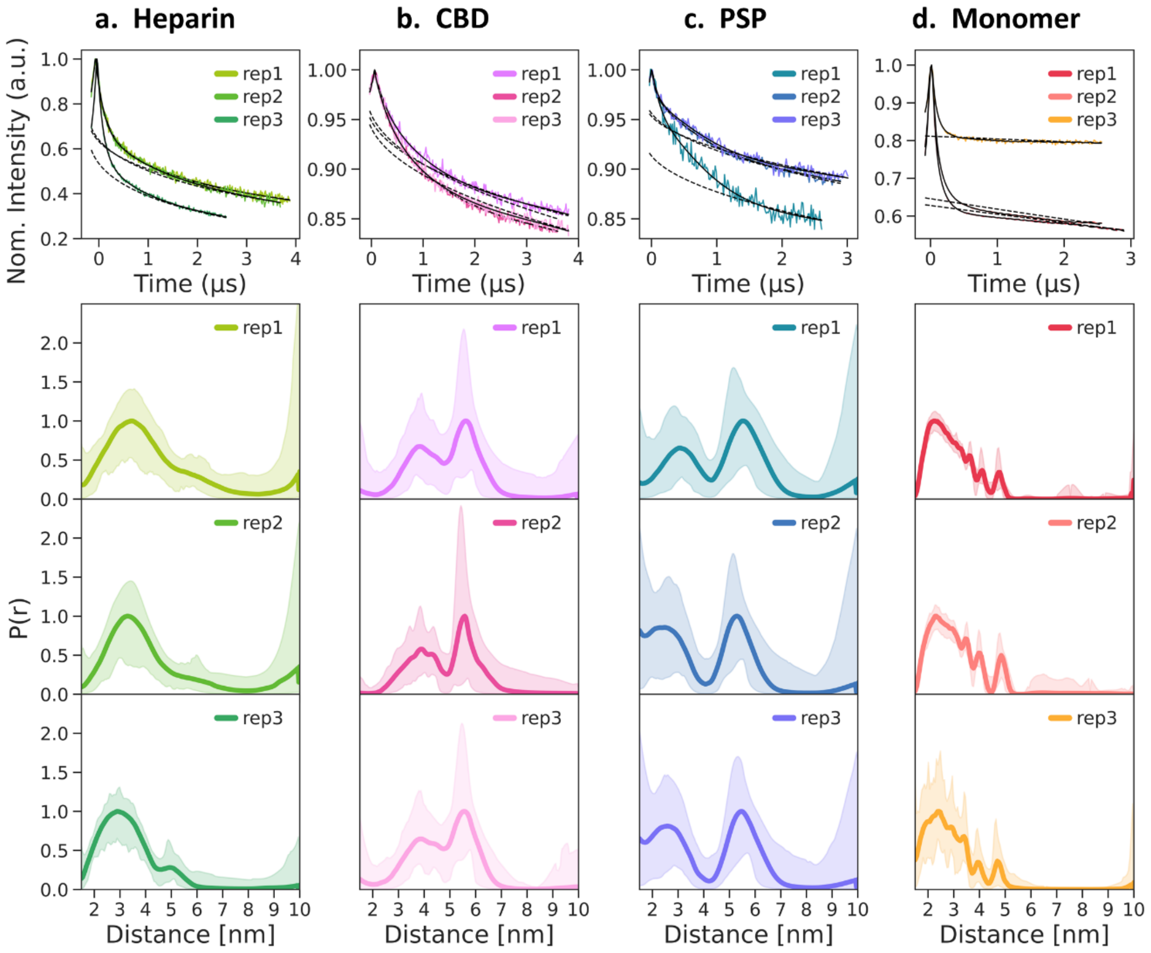


**Figure S5.** Technical replicates of P(r) measurements for residues 351 & 373. Independent triplicates (reps 1-3) of the intra-molecular DEER distance distribution, P(r), measured for residues 351 & 373 are shown for fibrils induced by (a) heparin, (b) cell-passaged CBD tau, (c) cell-passaged PSP tau, and (d) monomer. From top to bottom, figures show time-domain data with its fit and background fit, and distance distributions P(r).

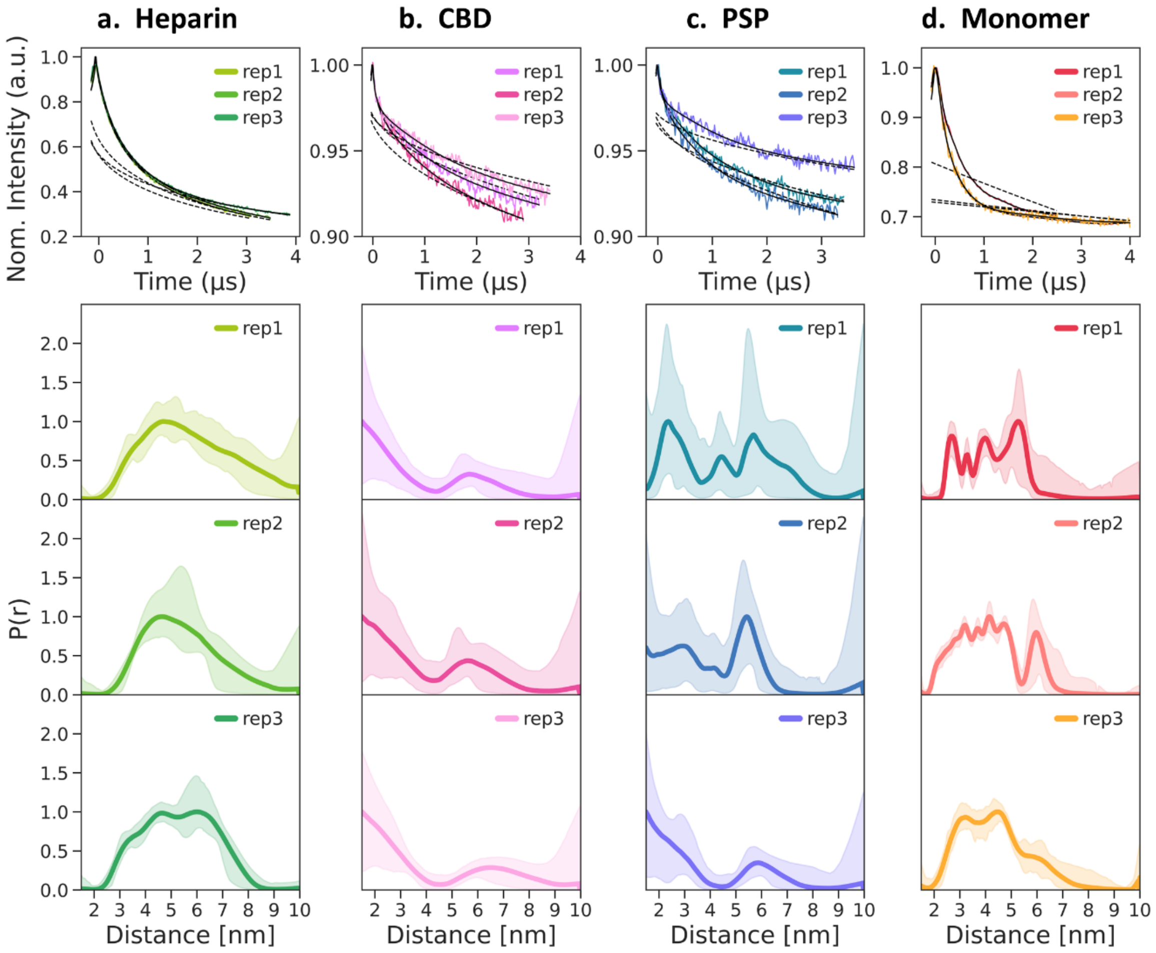


**Figure S6.** Technical replicates of P(r) measurements for residues 334 & 360. Independent triplicates (reps 1-3) of the intra-molecular DEER distance distribution, P(r), measured for residues 334 & 360 are shown for fibrils induced by (a) heparin, (b) cell-passaged CBD tau, (c) cell-passaged PSP tau, and (d) monomer. From top to bottom, figures show time-domain data with its fit and background fit, and distance distributions P(r).

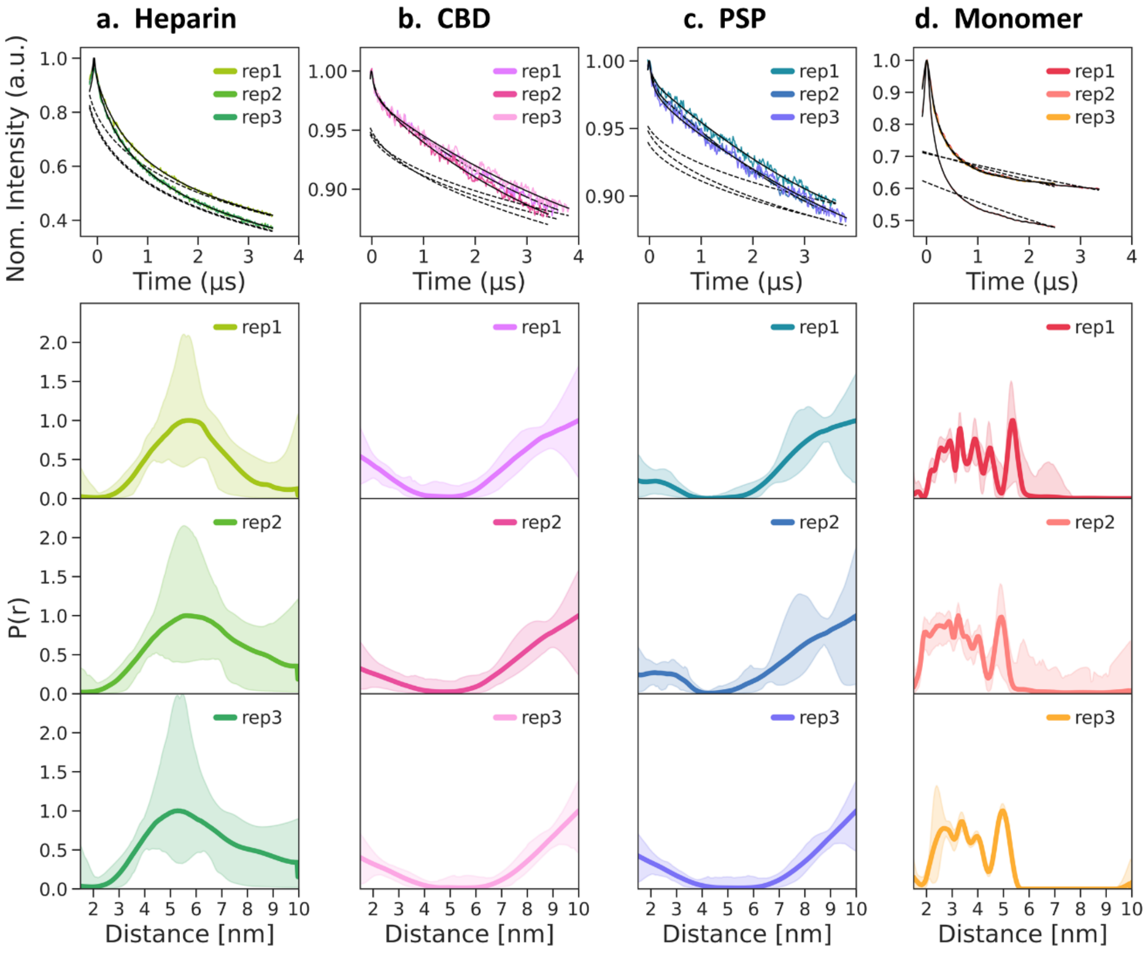


**Figure S7.** Technical replicates of P(r) measurements for residues 340 & 378. Independent triplicates (reps 1-3) of the intra-molecular DEER distance distribution, P(r), measured for residues 340 & 378 are shown for fibrils induced by (a) heparin, (b) cell-passaged CBD tau, (c) cell-passaged PSP tau, and (d) monomer. From top to bottom, figures show time-domain data with its fit and background fit, and distance distributions P(r).

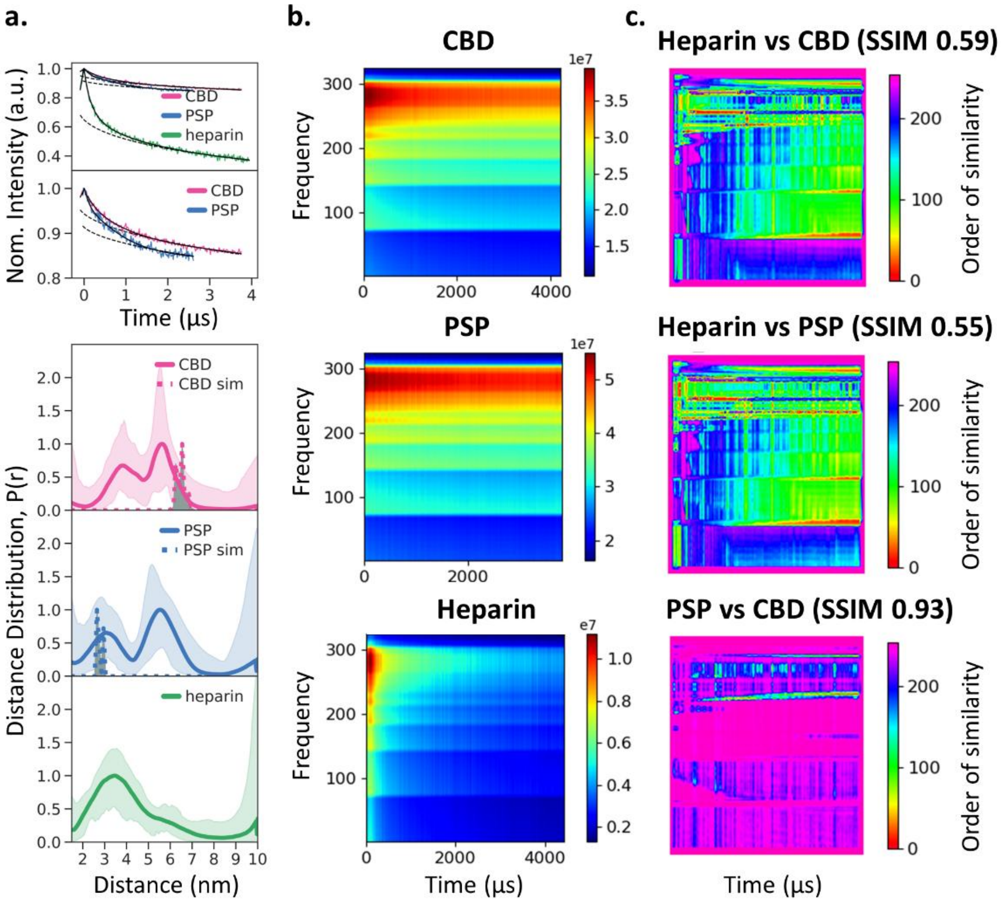


**Figure S8.** Pattern analysis of P(r) measurements for residues 351 & 373. (a) DEER time domain data (top) and P(r) (bottom) of Tau187 labeled at sites 351 & 373 are shown for cell-passaged CBD (pink), cell-passaged PSP (blue), and heparin seeded fibrils (green). (b) Spectrograms from CWT calculations of DEER time domain data are shown for cell-passaged CBD seeded fibrils (top), cell-passaged PSP seeded fibrils (middle), and heparin-induced fibrils (bottom). (c) Similarity gradient plots comparing spectrograms of heparin vs cell-passaged CBD fibrils (top), heparin vs cell-passaged PSP middle (middle), and cell-passaged PSP vs CBD seeded fibrils (bottom). The similarity gradient plot is shown as frequency vs time (top to bottom: high to low frequency bands) with the magnitude of similarity displayed in different colors, where red presents the least similarity and pink represents high similarity.

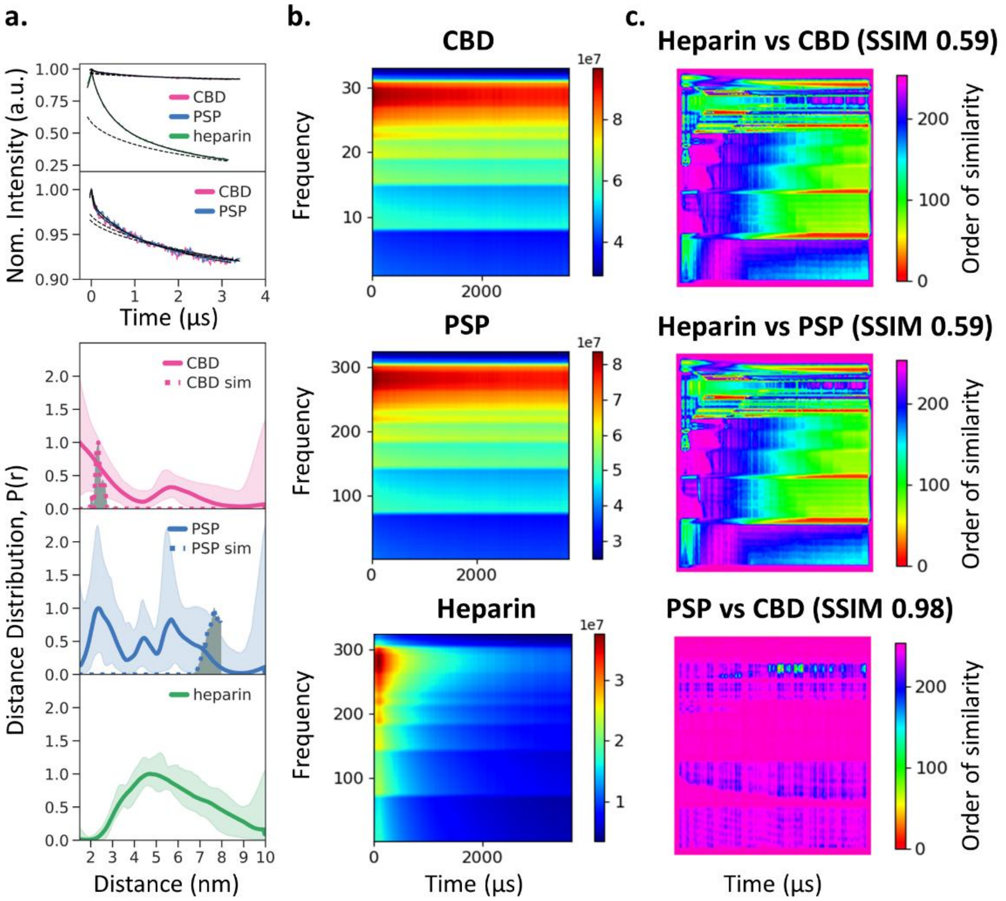


**Figure S9.** Pattern analysis of P(r) measurements for residues 334 & 360. (a) DEER time domain data (top) and P(r) (bottom) of tau187 labeled at sites 334 & 360 are shown for cell-passaged CBD (pink), cell-passaged PSP (blue), and heparin seeded fibrils (green). (b) Spectrograms from CWT calculations of DEER time domain data are shown for cell-passaged CBD seeded fibrils (top), cell-passaged PSP seeded fibrils (middle), and heparin-induced fibrils (bottom). (c) Similarity gradient plot comparing spectrograms of heparin vs cell-passaged CBD fibrils (top), heparin vs cell-passaged PSP middle (middle), and cell-passaged PSP vs CBD seeded fibrils (bottom). The similarity gradient plot is shown as frequency vs time (top to bottom: high to low frequency bands) with the magnitude of similarity displayed in different colors, where red presents the least similarity and pink represents high similarity.

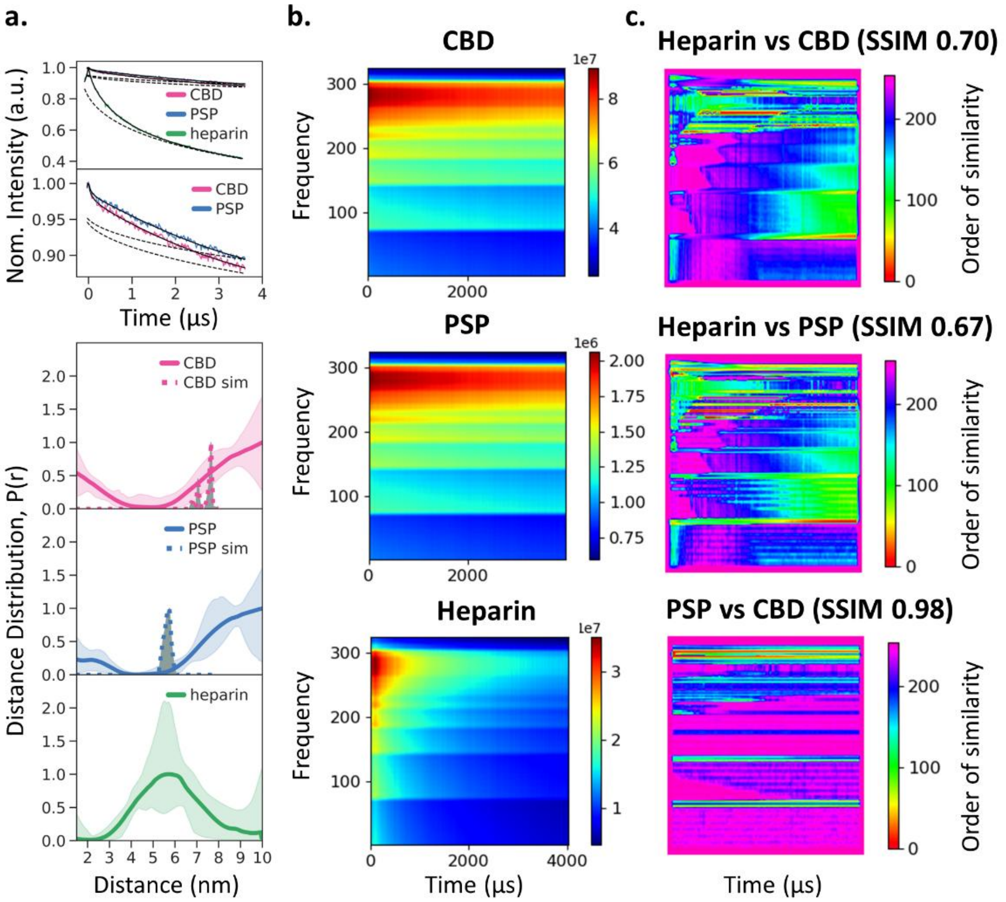


**Figure S10.** Pattern analysis of P(r) measurements for residues 340 & 378. (a) DEER time domain data (top) and P(r) (bottom) of tau187 labeled at sites 340 & 378 are shown for cell-passaged CBD (pink), cell-passaged PSP (blue), and heparin seeded fibrils (green). (b) Spectrograms from CWT calculations of DEER time domain data are shown for cell-passaged CBD seeded fibrils (top), cell-passaged PSP seeded fibrils (middle), and heparin-induced fibrils (bottom). (c) Similarity gradient plot comparing spectrograms of heparin vs cell-passaged CBD fibrils (top), heparin vs cell-passaged PSP middle (middle), and cell-passaged PSP vs CBD seeded fibrils (bottom). The similarity gradient plot is shown as frequency vs time (top to bottom: high to low frequency bands) with the magnitude of similarity displayed in different colors, where red presents the least similarity and pink represents high similarity.

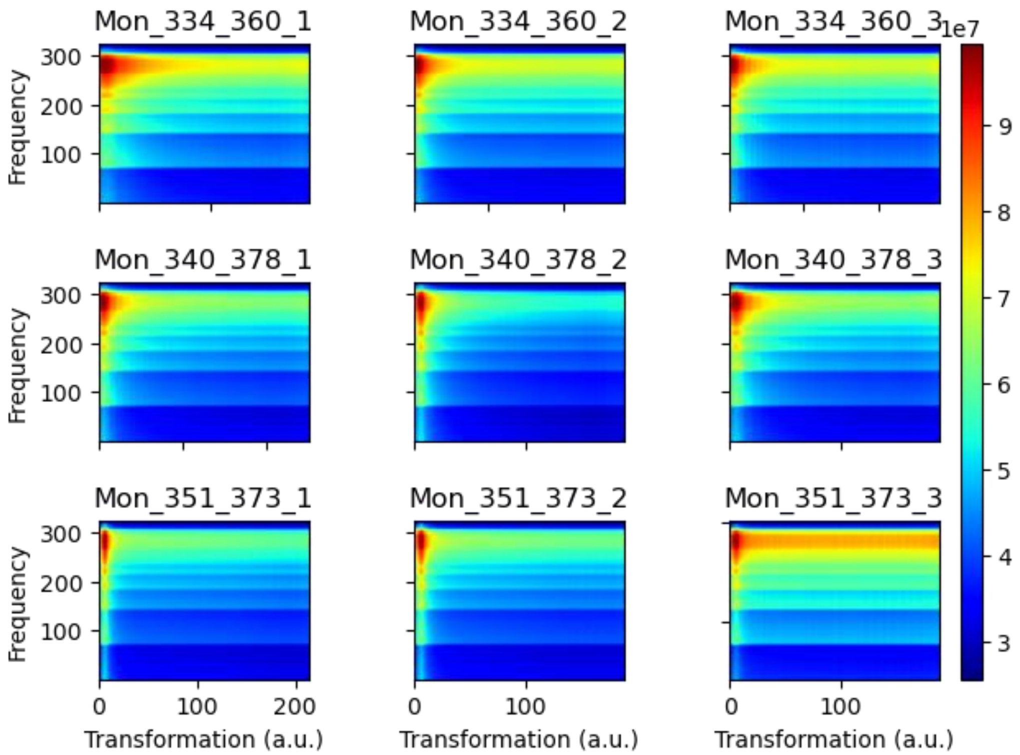


**Figure S11.** Spectrograms from CWT calculations of monomers. The spectrograms from CWT of independent triplicates (reps 1-3) of the DEER time-domain data of tau monomer labeled at residues 334 & 360 (top), 340 & 378 (middle), and 351 & 373 (bottom).

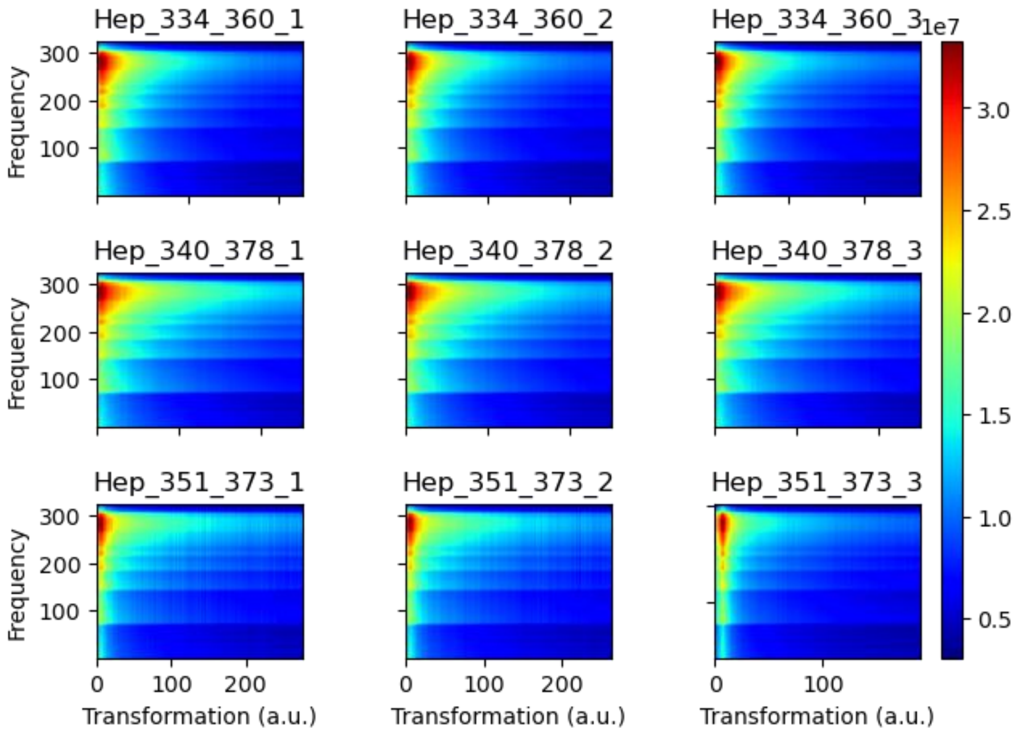

**Figure S12.** Spectrograms from CWT calculations of heparin induced fibrils. The spectrograms from CWT of independent triplicates (reps 1-3) of the DEER time-domain data of heparin-induced fibrils labeled at residues 334 & 360 (top), 340 & 378 (middle), and 351 & 373 (bottom).

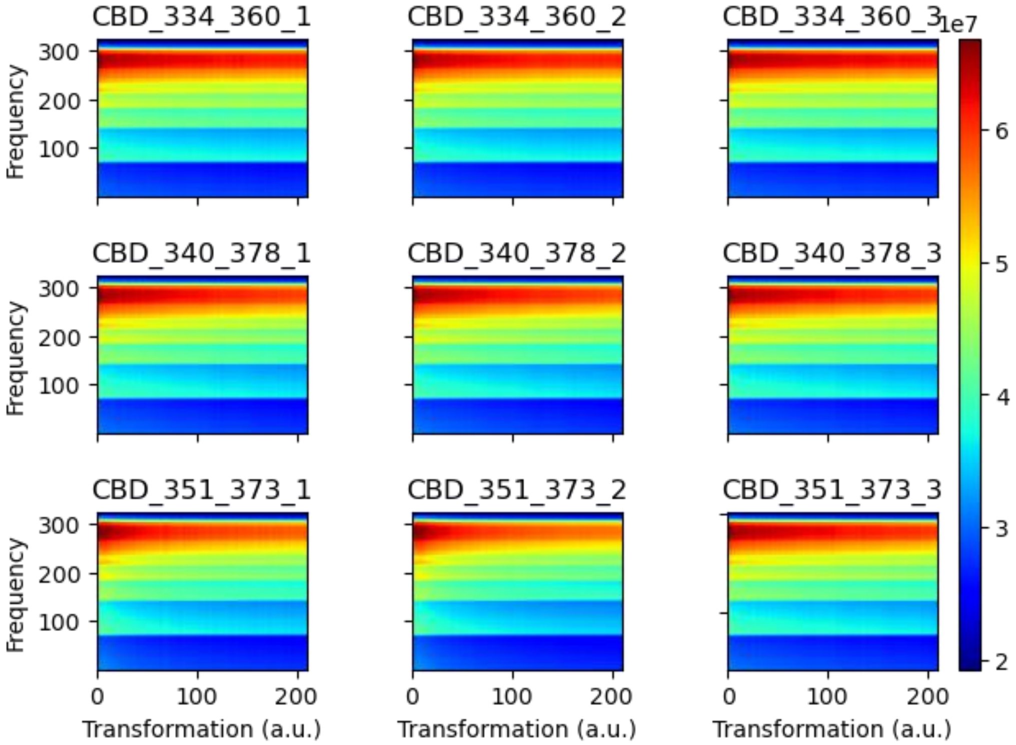


**Figure S13.** Spectrograms from CWT calculations of cell-passaged CBD seeded fibrils. The spectrograms from CWT of independent triplicates (reps 1-3) of the DEER time-domain data of cell-passaged CBD seeded fibrils labeled at residues 334 & 360 (top), 340 & 378 (middle), and 351 & 373 (bottom).

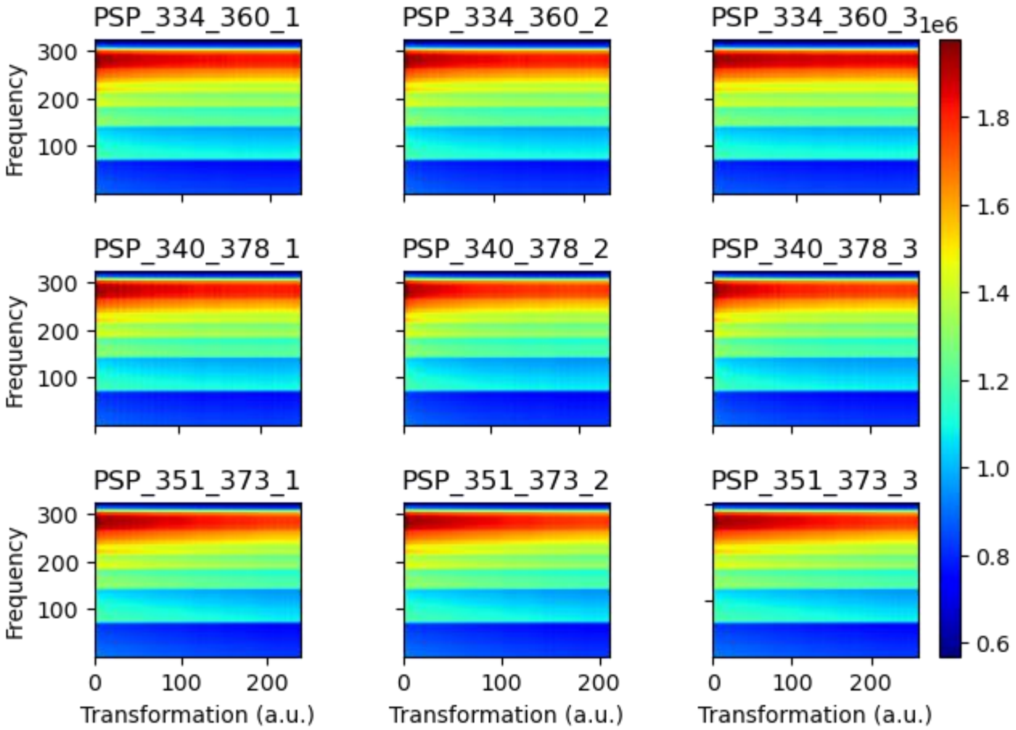


**Figure S14.** Spectrograms from CWT calculations of cell-passaged PSP seeded fibrils. The spectrograms from CWT of independent triplicates (reps 1-3) of the DEER time-domain data of cell-passaged PSP seeded fibrils labeled at residues 334 & 360 (top), 340 & 378 (middle), and 351 & 373 (bottom).

| Residues | Sample type | Replicate 1 | Replicate 2 | Replicate 3 | Avg SSIM | STD SSIM |
|---|---|---|---|---|---|---|
| 334-360 | Monomer | 0.837 | 0.853 | 0.841 | 0.844 | 0.007 |
| | Heparin | 1.000 | 0.963 | 0.963 | 0.975 | 0.018 |
| | CBD | 0.718 | 0.724 | 0.712 | 0.718 | 0.005 |
| | PSP | 0.727 | 0.729 | 0.714 | 0.723 | 0.006 |
| 340-378 | Monomer | 0.875 | 0.913 | 0.873 | 0.887 | 0.018 |
| | Heparin | 0.919 | 0.931 | 0.927 | 0.926 | 0.005 |
| | CBD | 0.722 | 0.724 | 0.718 | 0.722 | 0.002 |
| | PSP | 0.722 | 0.730 | 0.719 | 0.724 | 0.005 |
| 351-373 | Monomer | 0.862 | 0.866 | 0.788 | 0.839 | 0.036 |
| | Heparin | 0.885 | 0.900 | 0.921 | 0.902 | 0.015 |
| | CBD | 0.735 | 0.749 | 0.747 | 0.744 | 0.006 |
| | PSP | 0.720 | 0.733 | 0.730 | 0.728 | 0.005 |

**Table S1.** SSIM index of each spectrogram comparison to one set control spectrogram. Spectrograms for all DEER data were compared to that of the replicate one of heparin-induced fibril labeled at residues 334-360 (highlighted in yellow). The average and standard deviation of the replicates from each condition are displayed.
